# A Human Genome-wide CRISPR-Cas9 Screen Reveals Host Factors That Support Epstein-Barr Virus B cell infection

**DOI:** 10.64898/2026.09.28.754942

**Authors:** Hongbo Wang, Yao Yu Yeo, Chong Wang, Rui Guo, Shaowen White, Yifei Liao, Stephanie Yiu, Gloria Liu, Alvin Huang, David JS Zhao, Zongwei Wang, Sizun Jiang, Benjamin E. Gewurz, Bo Zhao

## Abstract

Epstein-Barr virus (EBV) infects 90% of the population worldwide and is associated with multiple malignancies, including Burkitt, Hodgkin and post-transplant lymphomas, nasopharyngeal and gastric carcinoma. EBV also triggers autoimmune diseases, including multiple sclerosis and systemic lupus erythematosus. While host factors important for EBV B-cell entry have been defined, including CD21/CR2, MHC class II and R9AP, much remains to be learned about host factors that support post-entry steps of EBV infection. To gain insight, we conducted a human genome-wide CRISPR-Cas9 screen. In addition to known EBV B cell receptors/co-receptors, we identified multiple new factors and pathways essential for EBV B cell infection. These included multiple B cell signaling pathways, actin cytoskeleton and nuclear import factors. Knockout of CD19, its tetraspanin chaperone CD81 or karyopherin subunit alpha 1 (KPNA1, also called importin-α5) significantly impaired establishment of EBV infection. EBV and CD19 co-localized at early stages of EBV infection, and the CD19 cytoplasmic tail was important for EBV infection, suggesting CD19 signaling supports EBV uptake. Depletion of KPNA1 significantly impaired establishment of EBV infection, but did not impair production of infectious EBV upon reactivation of latently infected B cells. Together, these results provide insights into pathways that support EBV B cell infection and identify potential therapeutic targets.

**Importance:** EBV infects ∼90% of humans and causes multiple cancers (Burkitt, Hodgkin, post-transplant lymphomas, nasopharyngeal and gastric carcinoma) and autoimmune diseases (MS, lupus). Yet while B cell entry receptors (CD21/CR2, MHC-II, R9AP) were known, the host machinery needed *after* entry — for the virus to actually establish infection was largely uncharacterized. This study fills that gap. Using CRISPR-Cas9 screening across the whole genome (rather than testing candidate genes), the study uncovered entire pathways B-cell signaling, actin cytoskeleton remodeling, and nuclear import as essential for infection, expanding the known host-dependency landscape well beyond entry receptors. Because these are discrete, druggable host factors tied to a specific stage of infection (establishment, not maintenance or reactivation), they represent candidate targets for preventing EBV-driven malignancies and autoimmune disease potentially more tractable than targeting the virus itself. The work moves EBV research beyond entry receptors to define the post-entry host dependencies that let the virus successfully infect B cells, identifying CD19 signaling and KPNA1-mediated nuclear import as new, targetable choke points relevant to EBV-associated cancers and autoimmunity.

## INTRODUCTION

The gamma-herpesvirus Epstein-Barr virus (EBV) persistently infects >90% of adults worldwide(1–3). Primary EBV infection typically occurs early in life and is often asymptomatic. However, when primary infection is delayed to late adolescence or early adulthood, it can cause infectious mononucleosis(2). EBV is etiologically associated with ∼350,000 malignancies per year(4). These include Burkitt lymphoma, Hodgkin lymphoma, central nervous system lymphomas, post-transplant lymphoproliferative disease, NK/T cell lymphoma, nasopharyngeal and gastric carcinoma(3). EBV infection increases multiple sclerosis risk by 32-fold (3, 5–10). EBV is also strongly linked to a wide range of other autoimmune diseases, including systemic lupus erythematosus (11), (12, 13). In EBV transformed B cells, EBV nuclear antigen 2 (EBNA2) DNA-binding sites are enriched for single-nucleotide polymorphisms linked to autoimmune disease susceptibility (14, 15). Therefore, there is growing interest in the development of vaccines and therapeutic approaches that limit B-cell infection. However, much remains to be learned about host factors that support early events in EBV uptake and the establishment of B-cell latency.

EBV infects both B and epithelial cells (16–18). EBV enters B cells through a well-characterized two-step process. First, EBV attaches to B cells through the interaction of EBV glycoprotein gp350/220 with CD21 (also called complement receptor 2 or CR2) or with CD35(19–21) . Then, EBV gp42 binds to the coreceptor, major histocompatibility complex (MHC) class II molecules (13–16), and triggers the core fusion machinery (glycoproteins H, L and B) to mediate host-virus membrane fusion and B cell entry (22–26) (16, 27–35).. R9AP has also recently been implicated as a co-receptor for EBV uptake through binding to EBV glycoproteins H and L (gH:gL)(36) . Viral capsids enter through EBV glycoprotein-driven endocytosis and fusion (8), which delivers them to the cytosol. Knowledge remains limited about how EBV capsids traffic to the nucleus to establish latent infection. EBV epithelial cell uptake is instead mediated by gH:gL, which bind to ephrin A2 (EphA2), DSC2, DSC3, and R9AP, and by gB, which binds to neuropilin 1 to trigger fusion at the plasma membrane (36–42). .

Following B-cell uptake, EBV tegument proteins and capsids are delivered to the cytosol(43) , (44), (45). There is limited knowledge of how EBV capsids traffic to the nuclear membrane. However, shortly after infection, EBV capsids dock at nuclear pore sites and the double-stranded, linear EBV genome is transferred to the nucleus(46) . Incoming EBV DNA is rapidly chromatinized and circularized. Rather than serving as the immediate template for EBV lytic replication, B cell infection typically results in EBV latency, in which a small number of EBV latency proteins and non-coding RNAs are expressed(47–49) (50) .

Despite well-established CD21 and MHC class II roles in support of EBV attachment at the earliest stages of B-cell entry, much remains unknown about subsequent events in EBV uptake and capsid trafficking. We therefore performed a human genome-wide CRISPR-Cas9 screen to identify host dependency factors important for early stages of EBV infection and latency establishment. Here, we highlight key roles for CD19 signaling and karyopherin-mediated nuclear import in EBV B cell infection.

## Results

### CRISPR-Cas9 screen for host factors that support EBV B cell infection

To identify host factors essential for the early stages of B-cell EBV uptake, we performed a genome-wide CRISPR-Cas9 screen in Daudi B cells with stable Cas9 expression. We chose these Burkitt cells for the screen, as they are highly infectable by EBV (40) and express a high level of CD21/CR2 **(Supplementary Fig.1a**). At a high EBV multiplicity of infection (MOI), we found that nearly 100% of Daudi cells could be super-infected by enhanced GFP (eGFP) encoding EBV, which enabled a screen for knockouts that impair EBV infection. As a positive control, we demonstrated that CRISPR-mediated CD21 depletion nearly abolished eGFP+ EBV infection (**Supplementary Fig.1a-b**).

We transduced 150 million Cas9+ Daudi cells with the Brunello pooled single guide RNA (sgRNA) CRISPR lentiviral library, which targets 19,114 human genes with 76,441 sgRNAs, typically four of which independently target each human gene(29). Daudi cells were infected at a MOI of 0.3 to minimize lentiviral co-infection. Successfully transduced cells were selected with puromycin three days post-transduction to generate a Daudi knockout pool, across which each human gene was targeted by CRISPR editing. Six days post-selection, 80 million successfully transduced, puromycin-resistant cells were harvested and stored at -80°C as the input control. 80 million cells were then infected with the eGFP expressing Akata EBV strain. To achieve near uniform B-cell infection, we used an EBV MOI of 10 Green Daudi Units (GDU, (51)). Three days post-infection, we used FACS to sort the 5% of cells with the lowest GFP signal, indicative of impaired EBV uptake or establishment of early latent infection **(Fig. 1a and Supplementary Fig. 1c-d).** Genomic DNA was purified from input versus sorted cells.

**Fig. 1.**
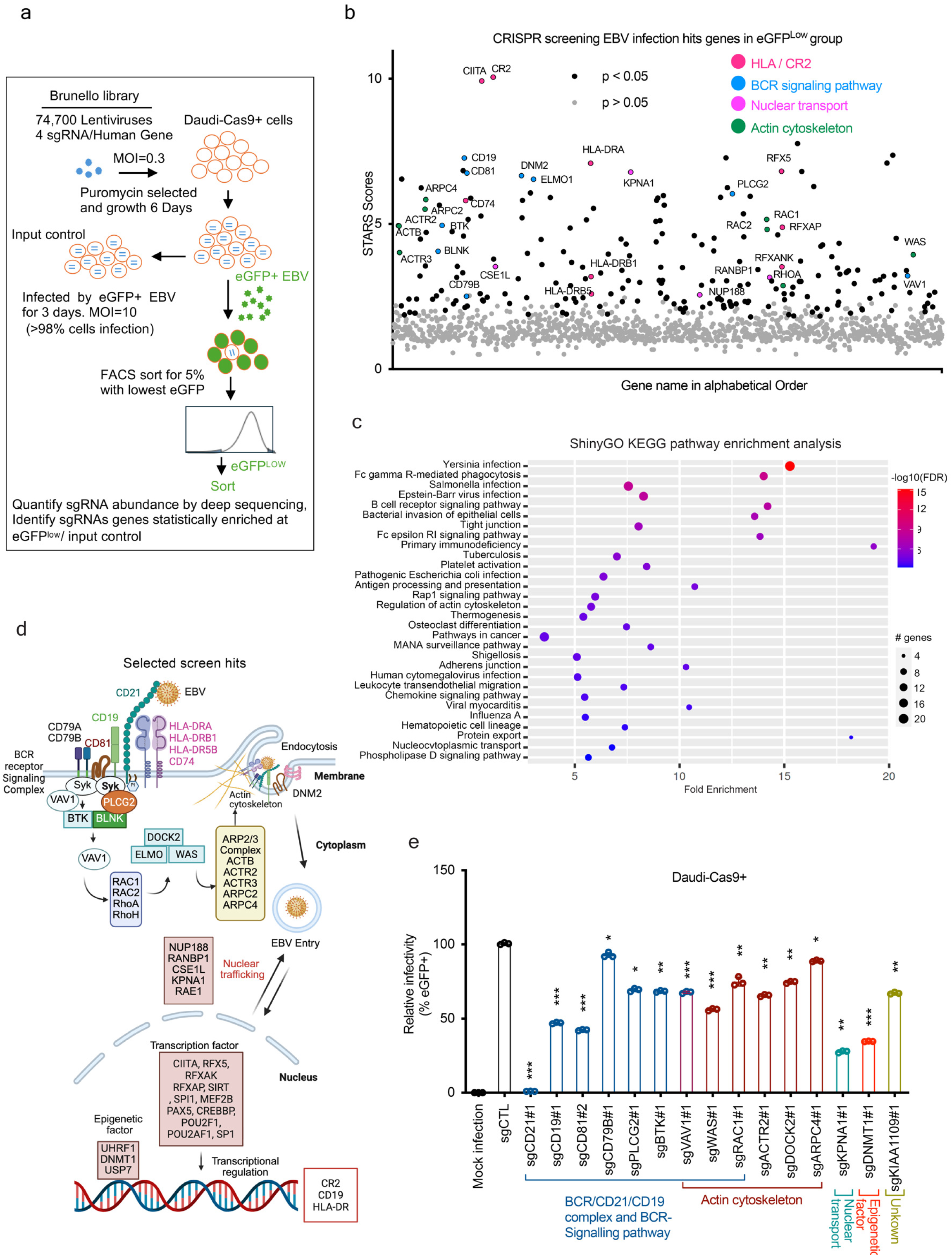
Human genome-wide CRISPR-Cas9 screen for host factors that support EBV B cell infection. a. Schematic diagram illustrating the CRISPR-Cas9 screen workflow. b. CRISPR screen hits. The y-axis shows the screen STARS score, a computational metric of screen signal. Higher STARS score indicates a stronger enrichment for sgRNAs targeting the indicated gene within the eGFP^low^ population versus the input control. Genes are ranked on the X-axis by alphabetical order. c. ShinyGO KEGG pathways enriched among screen hits at a p-value< 0.05 cutoff. d. Schematic highlighting selected CRISPR screen hits at a STARS scores > 2 and p-value< 0.05 cutoff, indicating proposed function in support of EBV infection. e. Validation of individual selected hits. Cas9+ Daudi B cells were transduced with either non-targeting sgRNA (sgCTL) or sgRNA targeting the indicated gene, infected with EBV at a MOI=2 for 3 hours, and then analyzed by FACS at two days later. The percentages of eGFP+ (EBV-infected) Daudi B cells were set to 100. Data represent mean ± SEM from three biological replicates. Two-sided unpaired Mann-Whitney test (***P < 0.001, **P<0.01 and *P<0.05).

Input versus sorted cell sgRNA abundance was quantified by PCR amplification and next-generation DNA sequencing. Signal-to-Noise Activity Ranking Score (STARS) analysis was used to identify screen hits, whose sgRNAs were enriched in the sorted, low-GFP cells relative to the input population. At an adjusted P-value<0.05 and STARS SCORE >2 cutoff(52), we identified 222 genes important for EBV B cell infection (**Supplementary Table 1-2**). Notably, top hits included CD21 and class II major histocompatibility complex trans-activator (CIITA), which is the master transcriptional coactivator that drives expression of MHC class II molecules **(Fig. 1b**) (**19, 22**). These results suggest that the CRISPR screen successfully identified host factors involved in EBV B cells infection. Additional top hits include factors involved in the B-cell receptor (BCR) pathway (CD79B), the B cell immunoglobulin co-receptor signaling pathways (CD19, CD81, SYK, BLNK and PLCG2), actin cytoskeleton regulation (actin, ARP2/3, ACTR3, ARPC2, and WAS), nuclear transport (KPNA1 and NUP188), and epigenetic regulation (DNMT1 and UHRF1), the latter of which are important for maintenance of EBV latency in Burkitt cells . The proposed biological functions of screen hits were analyzed by ShinyGO KEGG pathway enrichment analysis (**Fig. 1c-d**). Top pathways enriched amongst screen hits included EBV infection, B cell receptor signaling pathway, regulation of actin cytoskeleton and nucleocytoplasmic transport.

To further confirm these findings, we used CRISPR to deplete Daudi cell CD21. As expected, CD21 depletion nearly completely blocked infection by eGFP+ EBV **(Fig. 1e and Supplementary Fig. 1b)**. We also used CRISPR knockout (KO) to validate a range of additional top screen hits, including of the CD19 complex (CD19 or CD81), actin cytoskeleton regulators (VAV1, WAS1, ACTR2, ARPC4, and DOCK2), the nuclear karyopherin KPNA1 (also called importin-α5 or NPI-1), the DNA methylation enzyme DNMT1 and an uncharacterized gene KIAA1099. CRISPR KO of any of these screen hits significantly reduced eGFP+ EBV Daudi cell super-infection, supporting the roles of these genes and biological pathways in supporting EBV B cell infection **(Fig. 1e)**.

### CD19 intracellular signaling promotes EBV uptake

CD21, CD19, and CD81 form a complex that recognizes complement-tagged antigen. In this manner, complement-tagged antigens serve to co-localize B cell immunoglobulin and CD19/21/81 complexes(53) (54, 55). CD21 recognizes the complement tag, but has a short intracellular domain that is not thought to transmit intracellular signaling. Instead, interaction of the CD21 and CD19 extracellular domains facilitate signaling by the CD19 cytoplasmic tail, which contains YxxM/tyrosine motifs that recruit Src family kinases including Lyn to initiate downstream signaling(56){Tedder, 1997 #2621}. CD19 also recruits phosphoinositide 3-kinase (PI3K), the adaptor protein Vav, and other signaling molecules that regulate cytoskeletal rearrangement(56, 57). CD81 serves as a molecular chaperone to facilitate CD19 plasma membrane trafficking(56, 58) , though it is not required for CD21 cell surface trafficking(6, 59).

To further characterize their EBV uptake roles, we used CRISPR to deplete either CD19 or CD81 in Daudi and for cross-comparison also in EBV-negative Akata B cells (60). CD19 or CD81 depletion strongly impaired EBV super-infection of Daudi and of Akata cells. CD81 KO downregulated CD19 surface expression, but not *vice versa*, in both cell lines. Importantly, neither CD81 nor CD19 KO significantly affected plasma membrane expression of the EBV receptor CD21 or co-receptor MHC Class II (**Fig. 2a-b, Supplementary Fig. 2a-b**). These results suggests that CD19 and CD81 support EBV infection, likely downstream of EBV interaction with immunoglobulin receptor or MHC-class II co-receptor.

**Fig. 2.**
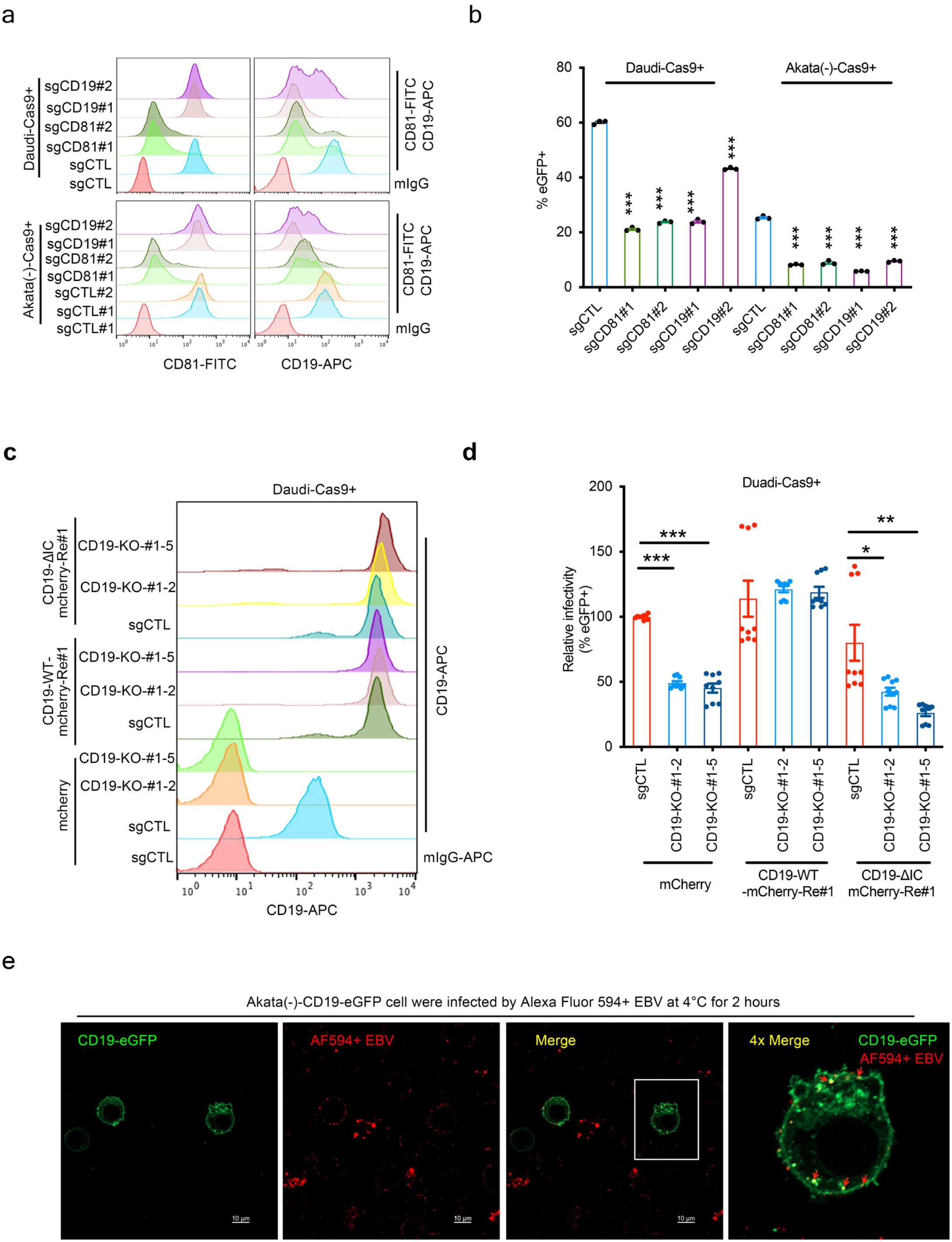
The plasma membrane proteins CD19 and CD81 support EBV B cell infection. a. Effects of CD19 and CD18 CRISPR KO on EBV B cell infection. Shown are representative FACS analysis of plasma membrane (PM) CD81 (left) or CD19 (right) expression in Cas9+ Daudi and EBV-negative Akata cells that expressed control sgRNA (sgCTL) or independent sgRNAs targeting CD19 or CD81. b. CD19 and CD81 KO effects on Daudi and EBV-negative Akata B cell infection. Shown are mean ± SEM percentages of eGFP+ (EBV-infected) Daudi (MOI=1) or Akata (MOI=10) cells from n=3 replicates. mIgG, mouse IgG negative control. c. Effects of Daudi CD19 KO and cDNA rescue on EBV uptake. Daudi CD19 KO single clones were transduced with lentiviruses expressing CRISPR resistant cDNAs encoding mCherry (control), mCherry-tagged CD19 wild type (WT, termed CD19-WT-mcherry-re#1) or mCherry-tagged CD19 intracellular domain deletion mutant (ΔIC, termed CD19-ΔIC mcherry-Re#1). PAM site mutations confer CD19 cDNA resistance to sgCD19#1 directed Cas9 editing. Two CD19 KO single cell clones (#1-2 and #1-5) were used. Shown are FACS analysis of plasma membrane CD19 expression. d. The CD19 intracellular domain supports Daudi B cell EBV uptake. Mean ± SEM percentages of eGFP+ Daudi cells, from n=9 replicates, using the cells shown in (c). Cells were co-incubated with eGFP+ EBV for 24 hours and then analyzed by FACS. The percentages of eGFP+ (EBV-infected) Daudi B cells expressing sgRNA control (sgCTL) and mCherry were set to 100. e. Confocal microscopy analysis of EBV and CD19 co-localization. Shown are representative confocal images of stably expressed CD19-eGFP and of Alexa Fluor 594 labeled EBV (AF594+ EBV). CD19 KO EBV-negative Akata cells were transduced with lentivirus that expressed an eGFP-CD19 cDNA rescue vector. Cells were co-incubated with Alexa Fluor 594-labelled EBV (AF594+ EBV) at 4°C for 2 hours. After one wash, CD19-eGFP and EBV co-localization was assessed. Enlarged images of the boxed areas are shown at right 4x the magnification. Scale bar: 10μm. Data represent mean ± SEM from three biological replicates. Two-sided unpaired Mann-Whitney test (***P < 0.001, **P<0.01 and *P<0.05).

We therefore hypothesized that the CD19 intracellular signaling supports EBV uptake, likely downstream of gp350 interaction with CD21. To test this, we used CRISPR to deplete endogenous CD19 and then express wildtpe or mutant CD19 rescue cDNAs. To do so, we first isolated CD19 KO Daudi single-cell clones. We confirmed that they were resistant to EBV infection **(Supplementary Figs. 2c-e),** but that plasma membrane levels of CD21, CD81 and HLA-DR (as a readout of MHC class II) remained comparable to parental Daudi cells **(Supplementary Fig. 2f)**. We then transduced CD19 KO cells with lentivirus encoding PAM-site mutated, CRISPR resistant CD19 rescue cDNAs, either wildtype CD19 (CD19-WT) or intracellular domain-deleted CD19 (CD19-ΔIC). CD19-WT and CD19-ΔIC were expressed at similar levels on the plasma membrane (**Fig. 2c**). WT CD19. However, whereas CD-19-WT (CD19-WT-Re#1) completely restored EBV infection, CD19-ΔIC (CD19-ΔIC-Re#1) failed to restore the infection. These results indicate that the CD19 intracellular signaling domain was essential for Daudi B cell EBV super-infection **(Fig. 2d and Supplementary Fig. 3)**.

We next examined if incoming EBV colocalizes with CD19 at the plasma membrane. To do so, we stably expressed eGFP-tagged CD19 in EBV-negative Akata CD19 KO cells, such that the CD19 pool was marked by GFP. The CD19-eGFP cells were co-incubated with EBV covalently labeled by the Alexa Fluor 594 (AF594) dye(61) at 4°C for 2 hours. AF594+ EBV particles frequently co-localized with CD19-GFP at the cell membrane, especially at regions with bright GFP+ spots that were presumably CD19 aggregates, suggestive of EBV-induced CD19 clustering (Fig. 2e).

In contrast to B-cell specific CD19, CD81 is more broadly expressed and for example is a co-receptor for hepatitis C virus (HCV)(62). We therefore next assessed if CD81 may have a direct role in promoting EBV infection, independently from chaperoning CD19. We used normal oral keratinocyte (NOK) epithelial cells(63) , as they are susceptible to EBV and model EBV oral infection. NOK cells express CD81 but are CD19-negative. We transduced Cas9+ NOK with lentivirus expressing control or *CD81* targeting sgRNA. Control versus CD81 depleted cells (**Fig. 3a**) were infected with EBV, either by cell-free virus or by cell-to-cell through co-culture with EBV+(eGFP+) Akata cells induced for lytic replication. Interestingly, CD81 KO increased EBV infection by cell free virus (**Fig. 3b-c**), but did not affect EBV uptake by the B cell-to-epithelial cell route (**Fig. 3d-e**). These data indicate suggest that CD81 itself is not required for EBV infection *per se*, though leave open the possibility that it may nonetheless play a CD19-independent role in B cell infection. Altogether, these results suggest that CD19 plays a key role in the earliest stages of EBV uptake, likely as part of the CD19/21/81 complex.

**Fig. 3.**
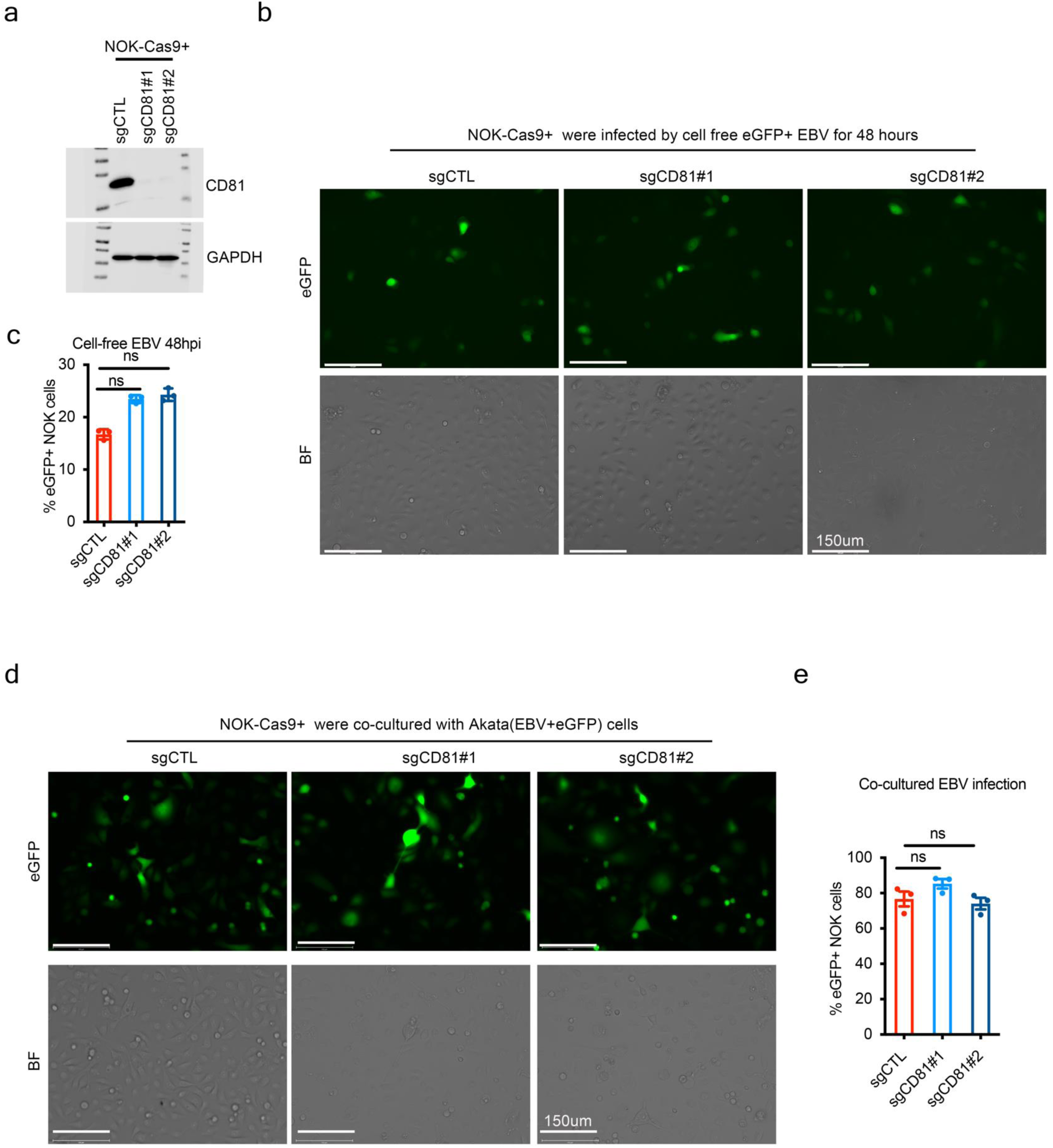
CD81-KO has no effect on EBV infection in normal oral keratinocytes (NOK). a. Immunoblot analysis of CD81 expression in whole cell extracts (WCE) from Cas9+ NOK cells that expressed control or CD81 targeting sgRNA. GAPDH served as a load control. b-c. CD81 is not required for NOK cells by cell-free EBV infection. FACS analysis of eGFP+ EBV infection of control (sgCTL) or CD81-KO (sgCD81) NOK epithelial cells. eGFP expression served as a reporter of EBV infection. eGFP positivity was quantitated by microscopy (b) or FACS (c) at 48h post-co-incubation. Scale bar: 150μm. d-e. CD81 is not required for Akata B cell to NOK epithelial cell EBV spread. FACS analysis of eGFP+ EBV infection of control or CD81-KO NOK cells. eGFP expression served as a reporter of EBV infection. eGFP positivity was quantitated by microscopy (d) or FACS (e) at 48h post-co-incubation. Scale bar: 150μm. Data represent means ± SEM from three biological replicates. two-sided unpaired Mann-Whitney test. ns, not significant.

### KPNA1 is required for EBV infection of B cells

Mechanisms underlying events immediately following EBV uptake, including as capsids transport to the nucleus, remain incompletely characterized. We therefore hypothesized that Karyopherin A1 (KPNA1) scored as a top screen hit (**Fig.1b**) due to obligatory roles in EBV capsid nuclear trafficking. In support, KPNA1 mediates nuclear import of multiple viral proteins, including human cytomegalovirus UL84 (64), HIV-1 Vpr (65), Ebola virus VP24 (66), and influenza virus polymerase (PB2) (67). Alternatively, it is not fully understood how EBV DNA passes through the nuclear pore(43), and an alternative hypothesis is that KPNA1 supports viral DNA transfer into the nucleus. A third hypothesis is that KPNA1 supports nuclear shuttling of a host protein important for EBV receptor expression or of an EBV protein important for the establishment of latency.

To distinguish between these, we first assessed if KPNA1 KO altered CD21, MHC class II, CD19 or CD81 plasma membrane expression. We transduced Cas9+ Daudi and EBV-negative Akata cells with lentiviruses that expressed control or KPNA1 targeting sgRNAs (**Fig. 4a**). In validation of the CRISPR screen result, KPNA1 depletion by either of two independent sgRNAs significantly reduced B cell EBV super-infection (**Fig. 4b-c and Supplementary Fig. 4a-c**). However, it did not affect plasma membrane expression of CD21, MHC class II (as judged by HLA-DR staining), CD19 or CD81 (**Fig. 4d**). We also generated Daudi control versus KPNA1 KO single cell KO clones. KPNA1 KO nearly completely suppressed EBV super-infection, as judged by the absence of EBV genomic eGFP expression (**Supplementary Fig. 4d-g**). However, once again, KPNA1 KO did not significantly alter expression of CD21, CD81, CD19 or the MHC class II HLA-DR allele (**Supplementary Fig. 4h**). To exclude off-target effects of CRISPR depletion, KPNA1 expression was rescued by cDNA with a PAM site mutation that engendered resistance to Cas9 cleavage (KPNA1-RE#1) (**Fig. 4e**). KPNA1 cDNA rescue restored successful EBV super-infection, as judged by eGFP expression (**Fig. 4f-g and Supplementary Fig. 5**). These findings support a key role for KPNA1 in promoting EBV infection in B cells at a level following EBV interaction with plasma membrane receptor.

**Fig. 4.**
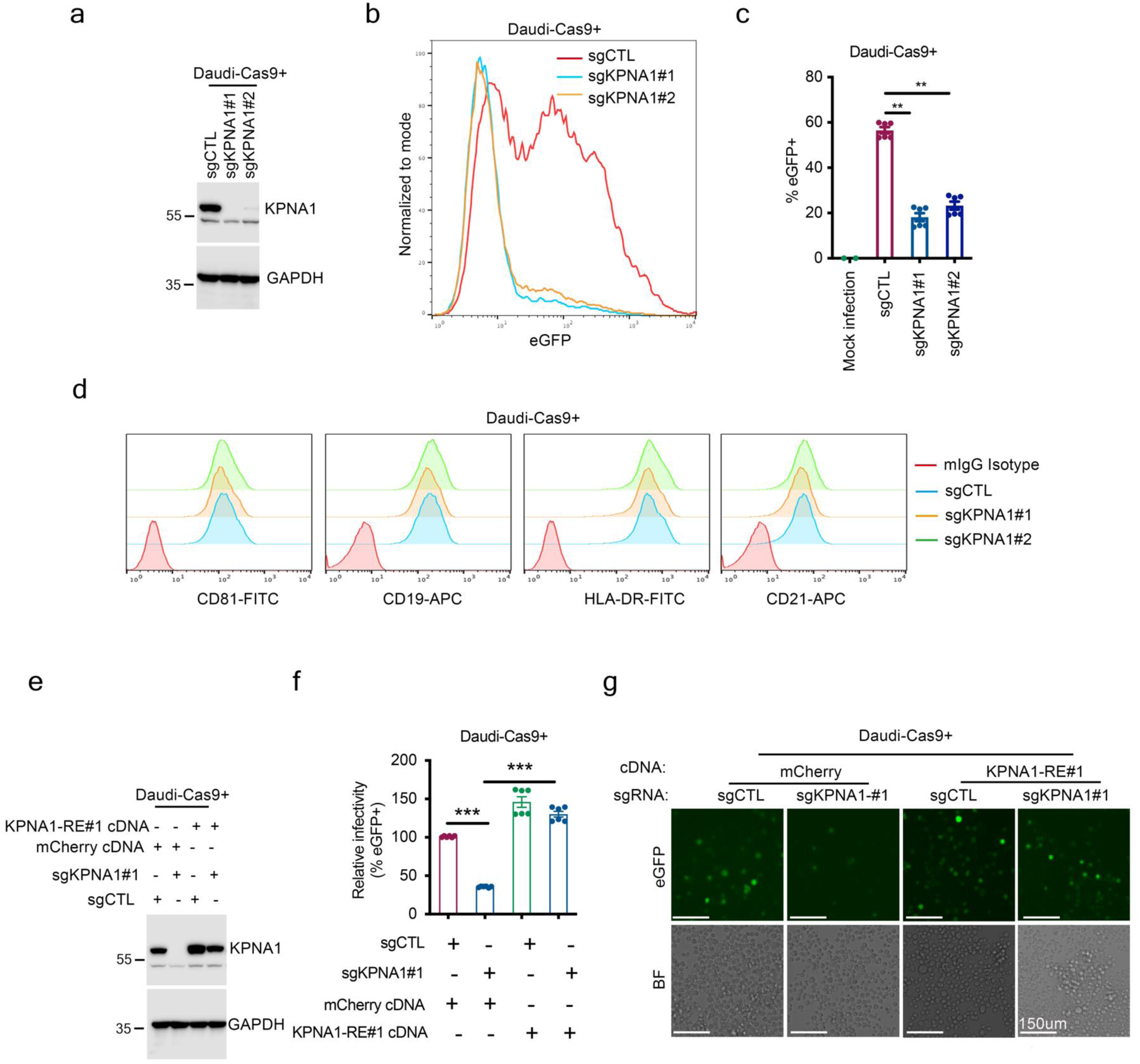
KPNA1 is required for EBV B cell infection. a. Immunoblot analysis of WCE from Daudi cells that expressed control or two independent CD81 targeting sgRNA. b-c. KPNA1 KO impaired EBV infection of Daudi B cells. Control or KPNA1 KO Daudi cells were co-incubated with EBV at a MOI of 1.0 for 3 hours at 37°C. The percentage of eGFP-positive infected cells was analyzed 24 hours post-infection using FACS. Representative histogram images of FACS were shown. c. Mean ± SEM % eGFP+ cells from n=3 replicates as in (b). d. The expression levels of PM CD81, CD19, HLA-DR and CD21 are similar in control versus KPNA1 KO Daudi cells. Cas9+ Daudi cells that expressed control or KPNA1 targeting sgRNAs were anaylzed by FACS. e. Analysis of KPNA1 expression in KPNA1-KO Daudi cells with KPNA1 cDNA rescue. Shown are immunoblots from WCL prepared from KPNA1-KO Daudi with stable mCherry (control) or CRISPR resistant KPNA1 (resistant to sgKPNA1#1) cDNAs. f-g. KPNA1 rescue restored EBV infection in Daudi-Cas9 cells. KPNA1-KO Daudi-Cas9 as in (e) were co-incubated with EBV at a MOI of 1.0 for 3 hours at 37°C. The percentages of eGFP-positive infected cells were analyzed 24 hours post-infection by FACS. The percentages of eGFP+ (EBV-infected) Daudi B cells expressing sgRNA control (sgCTL) and mCherry were set to 100. (g) Representative microscopy images of Cas9+ Daudi cells with the indicated sgRNA and rescue cDNA expression as in (e). Scale bar: 150μm. Data represent means ± SEM from three biological replicates. two-sided unpaired Mann-Whitney test (***P < 0.001 and **P<0.01). ns, not significant.

To further characterize the impact of KPNA1 depletion on B cell infection, we incubated EBV-negative Akata control versus KPNA1 KO cells with purified EBV at a MOI of 2 for 6 hours at 37° C. intracellular EBV DNA copy number was quantified by qPCR. KPNA1 depletion by either of two sgRNAs did not significantly alter EBV intracellular genome copy number (**Fig. 5a**). These results indicate that KPNA1 knockout does not impact EBV genome delivery into the B cell cytoplasm.

**Fig. 5.**
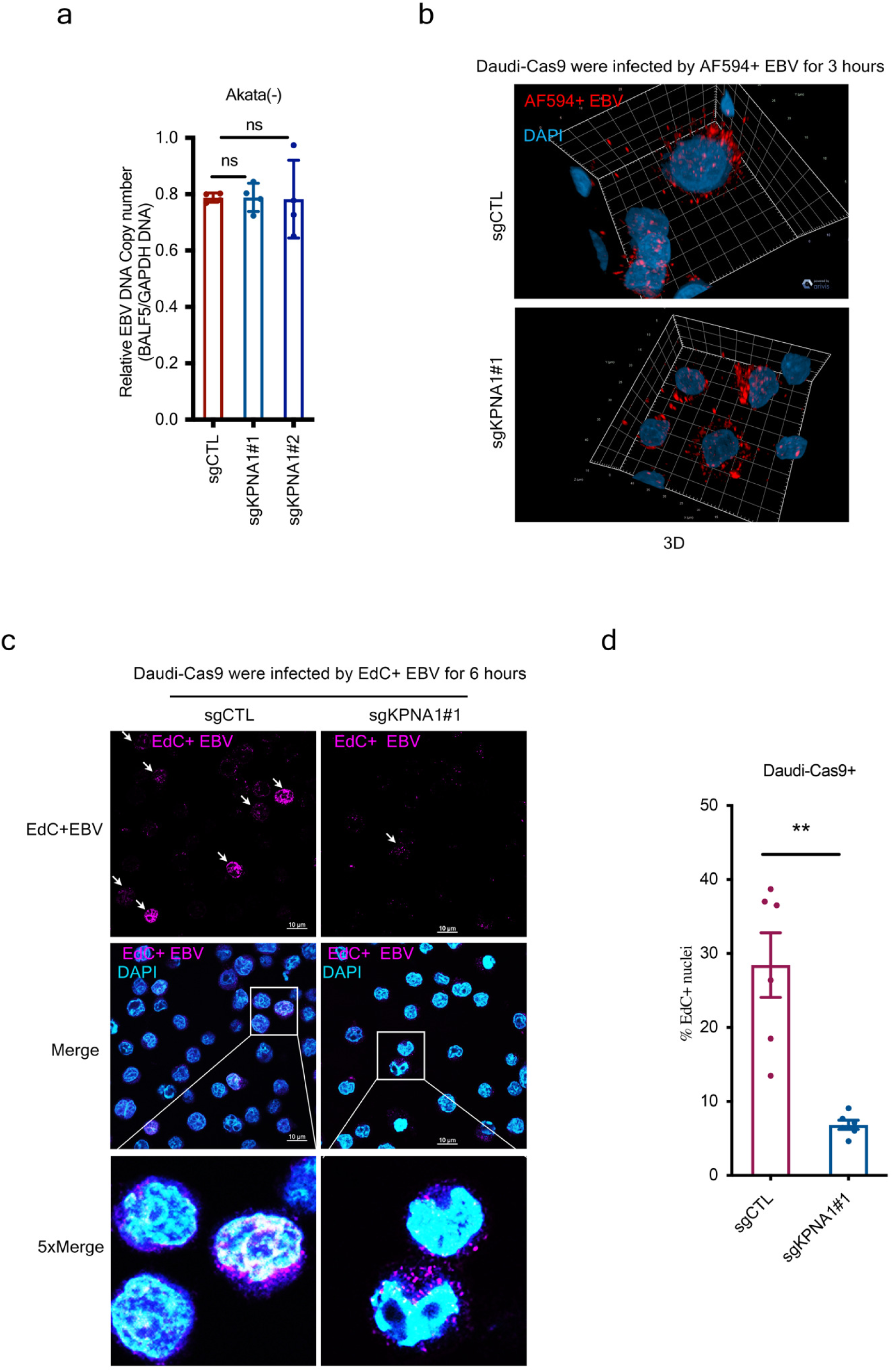
KPNA1 facilitates EBV genomic DNA nuclear transfer. a. KPNA1 is dispensable for EBV B cell infection. EBV-negative control or KPNA1 KO Akata B cells were co-incubated with EBV at a MOI of 2 for 6 hours at 37°C. After three washes with RPMI 1640, EBV genome copy # was determined by PCR quantification of the EBV BALF5 gene, normalized by GAPDH. EBV copy number was calculated as the ratio of EBV genome to GAPDH DNA copies. Shown are mean ± SEM from three biological replicates; ns, not significant. b. KPNA1 is dispensable for EBV entry in the B cell cytoplasm. Representative microscopy analysis of Cas9+ Daudi cells that expressed control or KPNA1 sgRNAs and co-cultured with Alexa Fluor 594-labelled EBV (Alexa Fluor 594+ EBV) for 3 hours at 37°C. After wash with RPMI 1640, cells were plated on slides and analyzed by 3D confocal microscopy. Alexa fluor 594+ EBV (red) and DAPI-stained nuclei (blue). c-d. KPNA1 KO impaired B cell EBV nuclear entry. Cas9+ Daudi control or KPNA1 KO cells were co-incubated with EdC-labelled EBV (EdC+ EBV) for 6 hours at 37°C, washed with RPMI 1640, plated on slides, processed by cycloaddition with azide-linked biotin and stained with avidin-Alexa Fluor 647. EBV genomic DNA was visualized using immunofluorescence confocal microscopy (EdC+ EBV purple, DAPI-stained nuclei blue). EBV genomic DNA abundance was quantified from n=6 images as in (c), using two replicates. Data represent means ± SEM from two biological replicates. two-sided unpaired Mann-Whitney test (**P<0.01). ns, not significant.

To next visualize KPNA1 KO impact on B cell infection, Cas9+ Daudi cells transfected with either control (sgCTL) or KPNA1 sgRNA and co-incubated with AF594-labeled EBV. We observed cytoplasmic viral particles in both control versus KPNA1 depleted cells by either 3D or 2D confocal microscopy (**Fig. 5b and Supplementary Fig. 6**). However, viral particles accumulated to a greater degree in KPNA1 KO cell cytoplasm (**Supplementary Fig. 6**). These results suggest that KPNA1 may play a key role following EBV B cell entry in capsid trafficking.

We next assessed if KPNA1 supports EBV genomic DNA nuclear transfer. To do so, we generated EdC-labeled genomes. Briefly, EBV+ Akata cells, which carried GFP-expressing EBV genomes, were induced for lytic reactivation by anti-IgG crosslinking in medium supplemented with EdC.

In this manner, EdC was incorporated into newly synthesized and packaged EBV genomes. EdC-labeled EBV was used to infect control or KPNA1 KO Daudi cells at 37°C for 6 hours, followed by the cycloaddition with azide-linked biotin. To visualize the EdC-labeled EBV DNA, cells were then stained with AF647-labeld avidin. Confocal microscopy revealed that KPNA1 KO strongly reduced EdC-labeled nuclear membrane and nuclear signal by comparison with control Daudi target cells **(Fig. 5c-d)**. Taken together, these data suggest that KPNA1 plays an important role in support of EBV genomic DNA nuclear entry.

### KPNA1 is not required for EBV epithelial cell infection

KPNA1 is expressed across a variety of cell types. To therefore evaluate if KPNA1 also promotes EBV epithelial cell infection, we tested KPNA1 KO effects on EBV NOK cells (**Fig. 6a**). NOK cells were co-incubated with 200-400 green Daudi units of EBV for 3 hours. FACS analysis revealed similar levels of EBV genomic driven GFP reporter signal in control and KPNA1 KO cells. Since the GFP reporter is expressed following EBV nuclear transfer, this result indicates that KPNA1 is not required for NOK EBV infection by cell free virus (**Fig. 6b-c**). Likewise, co-culture of Akata B cells triggered for lytic reactivation with control versus KPNA1 KO NOK cells resulted in similar levels of EBV genomic GFP reporter signal (**Fig. 6d-e**). These results suggest that different host factors, potentially distinct karyopherins, support EBV DNA nuclear transfer in B versus epithelial cells.

**Fig. 6.**
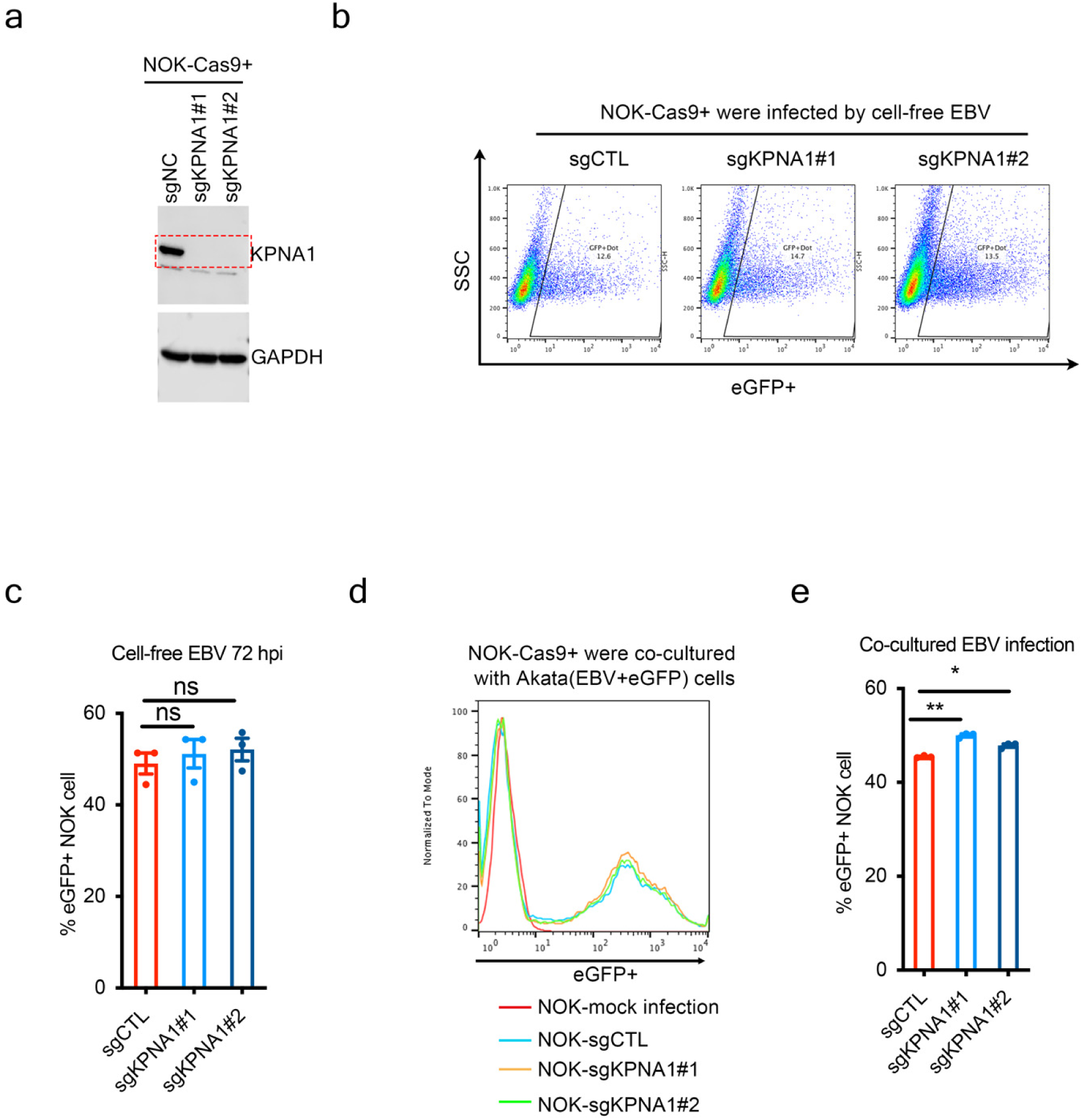
KPNA1 is not required for EBV entry into NOK cells. a. Immunoblot analysis of KPNA1 expression in WCL lysates of Cas9+ NOK with the indicated sgRNAs. b-c. KPNA1 is dispensable for NOK cell EBV infection. Control or KPNA1 KO NOK-Cas9+ cells were co-incubated with cell free EBV at a MOI of 200 - 400 Green Daudi Units (GDUs) for 3 hours at 37°C. eGFP+ cell percentages were quantitated by FACS 72 hours later. (b) Representative FACS images. (c) Mean ± SEM eGFP+ cell percentages from n=3 reps, as in (b). d-e. KPNA1 is dispensable for B cell to epithelial cell EBV spread. Control or KPNA1 KO NOK were cocultured with anti-human-IgG treated Akata-EBV+eGFP cells for 48 hours. NOK cell EBV infection was analyzed by FACS at 48 hours post-infection. (d) Representative FACS histograms. (e) Mean ± SEM NOK cell GFP+ cell percentages from n=3 reps, as in (d).

### KPNA1 is not required for the EBV B cell lytic cycle

Given its key roles in establishment of B cell infection, we next investigated whether KPNA1 also plays a role in EBV B cell lytic replication. To do so, we generated control versus KPNA1 KO EBV+ Akata cells. Cells were triggered for reactivation by anti-IgG crosslinking. Despite excellent KPNA1 depletion (**Fig. 7a**), we did not observe differences in the level of EBV reactivation, as judged by the EBV genomic GFP reporter signal, which is significantly brighter in reactivated cells (**Fig. 7b-c**). To further assess potential KPNA1 KO effects on EBV lytic replication, we tested infectious EBV levels in supernatants from control versus KPNA1 KO Akata cells stimulated by anti-IgG crosslinking. However, levels of infectious EBV were similar in control and KPNA1 KO supernatants, as judged by co-incubation with Daudi target cells (**Fig. 6d-**e). We also did not observe significant differences in cell-to-cell spread from control or KPNA1 KO Akata B cells to NOK epithelial cell targets (**Fig. 6f-g**). These data indicate that whereas KPNA1 plays a key role in B cell EBV genomic nuclear import, it does not play a substantial role in EBV lytic replication.

**Fig. 7.**
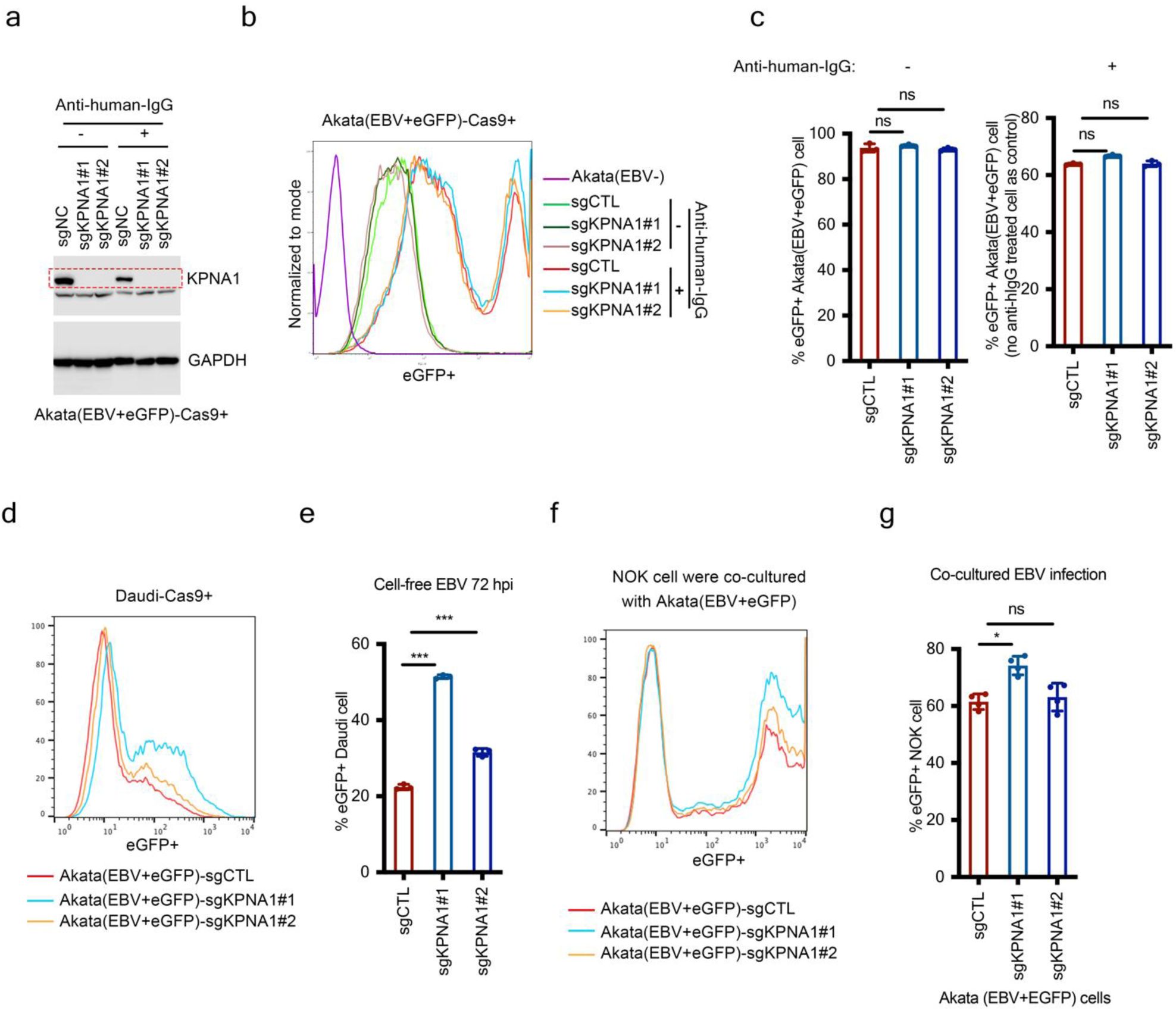
KPNA1 is dispensable for EBV replication in B cells. a. Immunoblot analysis of KPNA1 expression in WCL form Cas9+ Akata cells with control or KNPA1 sgRNAs, mock induced or induced for EBV reactivation by treatment with goat-anti-human-IgG for 24 hours. b-c. KPNA1-KO does not affect eGFP transgene expression in reactivated Akata (EBV+eGFP) cells. (b) Shown are representative FACS plots of GFP expression at 24 hours of mock treatment or goat-anti-human-IgG cross-linking. eGFP expression increases in lytic Akata EBV+eGFP cells. Representative histograms from n=3 replicates are shown. (c) Mean ± SEM eGFP+ cell percentages from n=3 replicates, as in (b). d-e. Akata (EBV+eGFP) cell KPNA1-KO does not affect EBV spread to B cell targets. Daudi B cell targets were co-incubated with supernatant from control or KPNA1 KO goat-anti-human-IgG cross-linked Akata (EBV+eGFP) cells, as in (a). Daudi eGFP expression was analyzed by FACS at 24 hours post-co-culture. Representative FACS histograms from n=3 replicates are shown. (d) Representative FACS histograms. (e) Mean ± SEM Daudi cell GFP+ cell percentages from n=3 reps, as in (d). f-g. Akata (EBV+eGFP) cell KPNA1-KO does not affect EBV spread to epithelial cell targets. NOK cell targets were co-incubated with supernatant from control or KPNA1 KO goat-anti-human-IgG cross-linked Akata (EBV+eGFP) cells, as in (a). NOK GFP expression was analyzed by FACS at 24 hours post-co-culture. Representative FACS histograms from n=4 replicates are shown. (f) Representative FACS histograms. (g) Mean ± SEM NOK cell GFP+ cell percentages from n=3 reps, as in (f). Data represent mean ± SEM from three biological replicates. Two-sided unpaired Mann-Whitney test ***P < 0.001 and *P<0.05). ns, not significant.

### KPNA1 does not induce innate immune responses during EBV infection

Karyopherins can support innate immune responses by supporting transcription factor nuclear entry, including of phospho-STAT1 (p-STAT1)(68, 69) , a key component of interferon (IFN) responses can inhibit viral replication and induce a cellular antiviral state. We therefore tested whether KPNA1 knockout alters infected B-cell interferon response, which could in turn potentially alter EBV infection. To do so, we used RT-qPCR to quantitate the levels of IFN-beta and interferon-induced gene OAS1 transcripts in control versus KPNA1 KO Daudi cells 24hours after co-incubation with EBV. However, EBV only weakly induced both IFN-beta and OAS1 expression and to a similar degree in control versus KPNA1 KO cells (**Fig. 8**). These data were also consistent with our ShinyGO KEGG enrichment analysis, which did not identify innate immune pathways as enriched amongst CRISPR hits (**Supplementary Table 2**). These results suggest that KPNA1 knockout did not score in our CRISPR screen as a result of effects on innate immune signaling, but rather because of its obligatory roles in nuclear EBV genome DNA transfer.

**Fig. 8.**
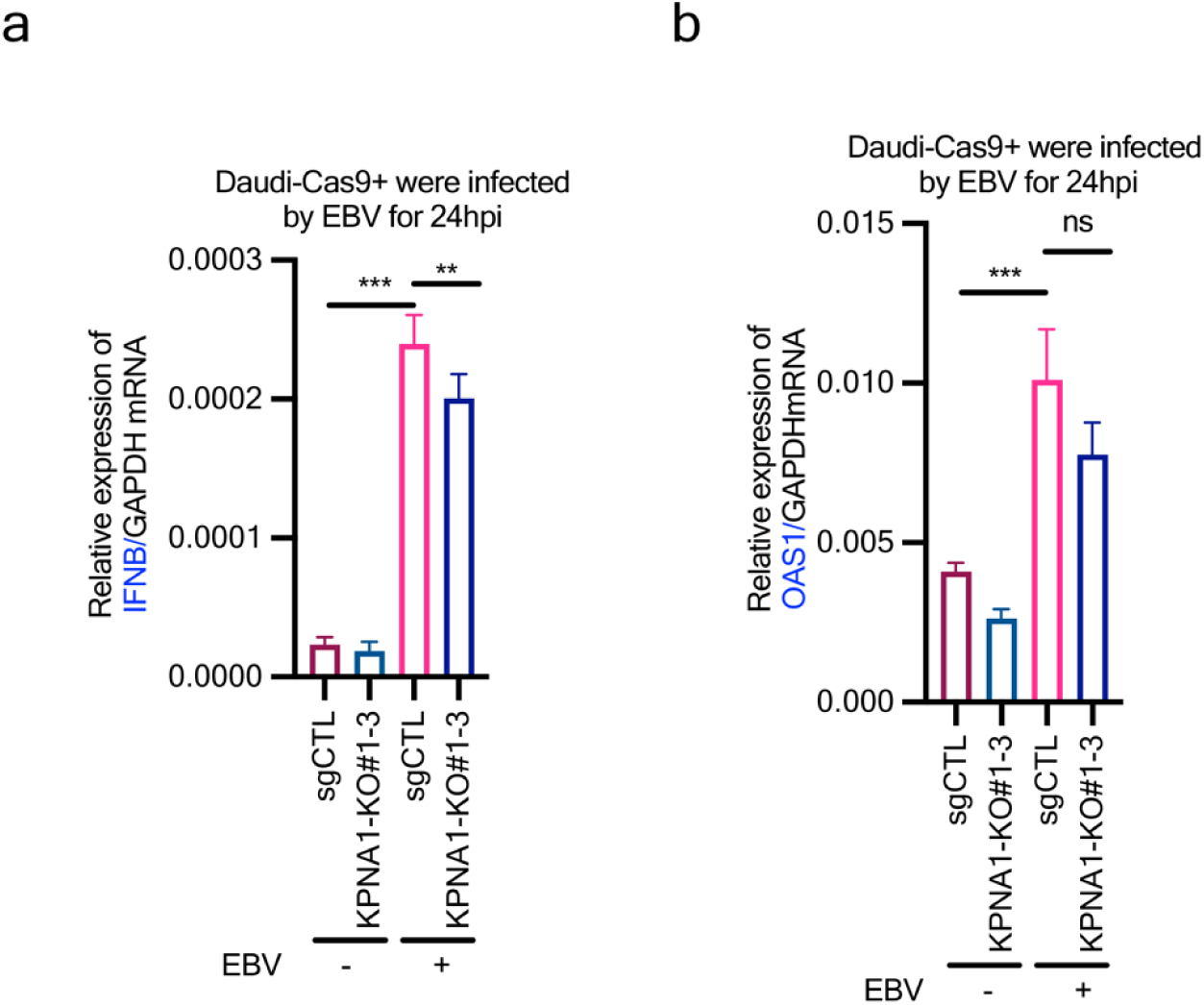
KPNA1 KO does not induce innate immune responses during EBV infection. a-b, KPNA1 KO does not promote IFNB or OAS1 mRNA expression in Daudi B cells. Control or KPNA1 KO Daudi cells were co-cultured with EBV at a MOI of 1.0 for 3 hours at 37°C. After 24 hours, mRNA levels of IFNB (a) and OAS1(b) were analyzed by real-time PCR. GAPDH was used as an internal control. Data represent mean ± SEM from three biological replicates. Two-sided unpaired Mann-Whitney test ***P < 0.001 and **P<0.01). ns, not significant.

## DISCUSSION

Although the principal EBV B cell attachment receptor CD21 and the gp42 MHC class II coreceptor have long been defined, host machinery that acts downstream of attachment to internalize incoming virions and to deliver EBV genomes to the nucleus has remained largely uncharacterized. Here, we used an unbiased genome-wide CRISPR-Cas9 screen in Daudi Burkitt B cells to systematically define host dependency factors for the establishment of EBV infection. The recovery of CD21 and of the MHC class II master transactivator CIITA validated the screen and provided a framework in which to interpret newly identified factors. Beyond these known receptors, the screen highlighted three interconnected modules: CD19/CD81 coreceptor signaling, actin cytoskeletal remodeling, and karyopherin-dependent nuclear import. We focused on CD19 and karyopherin subunit alpha 1 (KPNA1) as representative nodes acting at the plasma membrane and the nuclear envelope, respectively.

A notable implication of our screen is that it recovered the established EBV B cell receptors—CD21 and, through CIITA, MHC class II, but not apparently other new plasma membrane receptors. This result contrasts with analogous genome-wide CRISPR screens for epithelial cell entry factors, which identified desmocollin 2 (DSC2) as a principal EBV epithelial receptor that binds gH/gL, mediates membrane fusion, and functions upstream of EphA2(38) . The recovery of the known B cell receptors, in the absence of a comparable new receptor, suggests that the dominant plasma membrane determinants of EBV B cell tropism are now largely accounted for, and that the major remaining gaps concern post-attachment steps: coreceptor signaling, cytoskeletal remodeling, and nuclear delivery—rather than the identity of the attachment receptor itself.

CD19, CD21, and the tetraspanin CD81 assemble into the B cell coreceptor complex that lowers the threshold for antigen-driven B cell activation. Within this complex, CD21 recognizes complement C3d-opsonized antigen, but carries only a short cytoplasmic tail with no intrinsic signaling capacity. CD19 instead provides the signaling backbone through a long cytoplasmic tail whose tyrosine motifs, including YxxM motifs, recruit Lyn and other Src-family kinases, phosphoinositide 3-kinase (PI3K), Vav and phospho-PLCγ2. We propose that gp350-mediated CD21 attachment drives CD19 cytoplasmic tail phosphorylation and downstream signaling to support EBV uptake, likely through driving endocytosis, which enables subsequent EBV glycoprotein mediated fusion within endolysosomes.

Several observations support this model. First, a CD19 rescue cDNA lacking the intracellular domain trafficked normally to the plasma membrane, but failed to restore infection, indicating that CD19 signaling, rather than mere scaffolding, is required for EBV uptake. Second, incoming EBV colocalized with CD19 and appeared to induce CD19 clustering, as would be expected if virion-crosslinked receptor nucleates signaling microdomains. Third, the screen independently recovered the proximal signaling effectors SYK, BLNK, and PLCG2, as well as the actin-regulatory machinery, including VAV1, WAS, Arp2/3 subunits (ARPC2, ARPC4, ACTR2, ACTR3) and the guanine nucleotide exchange factor DOCK2. CD19-nucleated PI3K and Vav signaling activates Rac-family GTPases and WAVE/WASP/Arp2/3 to drive branched actin polymerization, a well-described requirement for receptor-mediated endocytosis. We therefore favor a model in which gp350-CD21 attachment triggers CD19 signaling, which in turn reorganizes cortical actin to internalize the attached virion, thereby coupling attachment to productive uptake.

The role of CD81 in our system appears to be principally to support CD19 plasma membrane expression, rather than to serve as an independent EBV entry factor. In CD19-negative NOK epithelial cells, CD81 depletion did not impair cell-free EBV infection, and it had no effect on B cell-to-epithelial cell transmission. These findings argue that CD81 is not itself an obligatory EBV receptor, and are consistent with its broader roles as a tetraspanin scaffold. They nonetheless leave open the possibility that CD81 contribute to the organization of the CD19 complex.

The identification of the karyopherin-α family member KPNA1 as a host dependency factor points to a discrete and previously underappreciated step in EBV B cell infection, namely import of the incoming viral genome across the nuclear envelope. We suspect that the karyopherin/importin beta subunit did not score in our screen as its KO may have been lethal. For instance, KPNB1 is the shared translocation receptor for hundreds of NLS cargoes. KPNA1 depletion did not alter surface expression of CD21, MHC class II, CD19 or CD81, and it also did not reduce the EBV genome copy number delivered to the cytoplasm, indicating that attachment, fusion, and cytoplasmic entry proceed normally in its absence. Rather, KPNA1 loss caused cytoplasmic accumulation of incoming EBV capsids, together with a marked reduction in nuclear and perinuclear signal from EdC-labeled viral genomes, localizing the defect to nuclear genome delivery. These observations place EBV alongside other DNA viruses that co-opt the classical importin-α/importin-β nuclear import pathway. For the prototypical alphaherpesvirus herpes simplex virus 1 (HSV-1), incoming capsids traffic along microtubules by dynein and dynactin to the nuclear pore complex (NPC), dock at the cytoplasmic face of the NPC through interactions involving importin-β and the nucleoporins Nup214/CAN and Nup358/RanBP2, and eject their genome through the pore. The HSV-1 inner tegument protein pUL36 (VP1/2) contributes a nuclear localization signal important for capsid–NPC docking and genome release .

Karyopherin-α proteins serve as the adaptors that bridge classical NLS-bearing cargo to importin-β and the Ran-GTP-powered translocation machinery. They mediate nuclear import of numerous viral proteins, including human cytomegalovirus UL84, HIV-1 Vpr, Ebola virus VP24, and influenza virus PB2. By analogy, KPNA1 may recognize an NLS within an EBV capsid or tegument component, plausibly a portal-associated or capsid vertex-specific protein, or a major tegument protein such as BNRF1. In this manner, it may dock incoming EBV capsids at the NPC and license genome ejection. Alternatively, KPNA1 may instead act more directly to facilitate EBV genome translocation, a step whose mechanism remains incompletely defined.

Karyopherin contribution to the delivery of the incoming herpesvirus genomes has been dissected in greatest mechanistic detail for HSV-1. Following plasma membrane fusion, the HSV-1 nucleocapsid is transported along microtubules to the nuclear periphery, docks at the NPC, and injects its DNA into the nucleoplasm(8–10, 70) (7). Cell-free reconstitution of this step established that capsid attachment to the NPC and subsequent genome uncoating depend on the classical nuclear-import machinery. Importin-β (KPNB1) is both necessary and sufficient to support capsid binding to isolated nuclei, whereas efficient DNA release also required cytosol as well as metabolic energy (7). However, KPNA1 was dispensable for docking, and the Ran-GTP/GDP cycle was required. These findings suggest that the HSV-1 capsid engages KPNB1 and the FG-nucleoporin meshwork, much as a conventional import substrate does, rather than through a KPNA1-adaptor-mediated NLS interaction. Consistent with a direct capsid-pore interface, the capsid-associated large tegument protein pUL36 (VP1-2) and the minor capsid protein pUL25 are the principal viral determinants of docking, and their engagement has been linked to specific nucleoporins, namely Nup358/RanBP2(10) and Nup214/CAN, together with hCG1(8) , which redistribute during entry.

The capsid-side determinant that couples this process to the import system is a short, highly basic nuclear localization sequence (NLS) in the HSV-1 VP1-2/pUL36 N-terminal region. This motif functions as an efficient NLS in isolation(71) , and a deletion mutant that perturbs this sequence arrests HSV-1 infection at or immediately upstream of NPC docking(72) . Similar NLS elements are present in the VP1-2 homologues of all three herpesvirus subfamilies, arguing that importin-dependent capsid targeting is a shared strategy across the alpha-, beta-, and gammaherpesviruses (73).

With regards to herpesvirus DNA transfer, the karyopherins appear to mediate docking and licensing of the pore, rather than actively translocating the genome, whose ejection is driven substantially by the internal pressure of the packaged DNA(74, 75) . Furthermore, KPNA1 involvement is required for the nuclear import of HSV-1 regulatory and replication proteins and for efficient capsid assembly, but appears to be dispensable for nuclear targeting of the incoming HSV-1 capsid itself (76). Direct biochemical dissection of these interactions has remained largely confined to HSV-1, and the importin dependence has therefore been inferred for CMV, VZV, and the gamma-herpesviruses.

Several features of the EBV KPNA1 requirement are noteworthy. First, KPNA1 was dispensable for EBV infection of NOK epithelial cells, implying that a distinct karyopherin-α paralog supports nuclear genome delivery in epithelial cells. This finding is consistent with the well-documented cell-type and cargo-specific importin-α family member division of labor(77) (78). Second, KPNA1 was not required for EBV lytic replication, infectious virion production, or cell-to-cell spread from reactivated B cells, indicating that the nuclear import demands of an incoming genome differ from those of *de novo* lytic DNA replication and nuclear egress, the latter of which proceeds through the EBV nuclear egress complex(79, 80) Finally, EBV only weakly induced IFN-β and OAS1, to comparable degrees in control and KPNA1-depleted cells, arguing that KPNA1 scored in the screen because of an obligatory role in nuclear genome delivery, rather than through effects on karyopherin-dependent interferon signaling.

Taken together, our results define a route by which EBV converts attachment into productive nuclear infection. gp350-CD21 engagement triggers CD19 coreceptor signaling and actin remodeling to internalize the virion, after which KPNA1-dependent nuclear import delivers the genome for chromatinization and the establishment of latency. This route reinforces the theme that EBV is intimately adapted to the B cell immunoglobulin and coreceptor signaling platform at multiple stages of its life cycle, exploiting CD19 coreceptor signaling to gain entry and, upon reactivation, deploying BALF0/1 to degrade the B cell receptor and immunoglobulin in support of virion egress(81) . The dependency factors identified here suggest candidate points of therapeutic intervention, as agents that interrupt CD19-nucleated signaling or karyopherin-α-dependent nuclear import could in principle limit the establishment of EBV infection. Such approaches would, however, need to contend with the essential physiological roles of CD19 signaling in B cell development and of KPNA1 in host nucleocytoplasmic transport.

Several limitations of our study warrant discussion. The screen was performed by superinfection of the Daudi Burkitt cell line, which expresses high levels of CD21 and is highly infectable. Although this facilitated a robust, near-uniform infection readout, it does not fully recapitulate the resting primary B cell, and the reliance on an EBV genomic GFP reporter as the readout for the establishment of infection means that hits could in principle act at attachment, uptake, nuclear delivery, or early latency gene expression. For KPNA1, our data establish a requirement for nuclear genome delivery, but do not yet distinguish whether KPNA1 acts directly, by importing a capsid- or genome-associated NLS cargo, or indirectly, and the specific EBV protein recognized by KPNA1 remains to be defined. Similarly, while the coordinate recovery of CD19, its proximal signaling effectors, and the actin machinery strongly implicates CD19-driven signaling in virion uptake, the individual contributions of each cytoskeletal and signaling hit, and the precise endocytic route taken by internalized virions, will require further dissection. Extending these findings to primary B cells will be an important goal for future work.

## Methods

### Reagents

The reagents used were as follows: antibodies against KPNA1 (sc-101292, Santa Cruz), CD19-APC (302212, Biolegend); CD81-FITC (349504, Biolegend); CD81(52892S, CST); CD21-APC (354906, Biolegend); HLA-DR-FITC (327006, Biolegend); GAPDH (sc-47724 HRP, Santa Cruz); Anti-V5 (13202S, CST); HA (ab9110, abcam); HA Magnetic Beads (88837, Invitrogen); Anti-rabbit IgG, HRP-linked Antibody (#7074, CST); Anti-mouse IgG, HRP-linked Antibody(#7076, CST); Polybrene (TR1003G, sigma); anti-human IgG (A0423, DAKO); EdC (T511307, Sigma); Alexa Fluor™647-advin (S21374, Invitrogen™); TransIT®-LT1 Transfection Reagent (MIR 2306 , Mirus Bio). Opti-MEM™ (11058021, Invitrogen™); ProLong Diamond Antifade with DAPI (P36935, Invitrogen™). All other reagents were obtained from Sigma-Aldrich, unless otherwise indicated.

### Cell culture

Akata EBV-negative, Daudi and BJAB are Burkitt lymphoma cell lines. Akata (EBV+eGFP) is a cell line that produces EBV with the eGFP cDNA integrated into the BXLF1 gene locus. In this manner, EBV expresses the eGFP protein in infected cells. These cell lines were cultured in RPMI 1640 medium (GIBCO) supplemented with 10% fetal bovine serum (FBS, GIBCO), 1% streptomycin and 1% penicillin. HEK-293T cells were cultured in Dulbecco’s modified Eagle’s medium (DMEM, Life Technologies) with 10% FBS, 1% streptomycin and 1% penicillin. All cells were grown in a humidified 5% CO2 incubator at 37°C and passaged using standard cell culture techniques.

### Vectors

To generate the expression vectors for CD19, CD19 cDNA was PCR-amplified from a commercially available plasmid, pLenti-CD19-V5 (HsCD00436263, DNASU), and integrated into either pCDNA3.1-eGFP or into pCDNA3.1-mCherry. A CD19 cDNA mutant with PAM sequence resistant to CD19 #1 sgRNA was generated from pCDNA3.1-CD19-eGFP and pCDNA3.1-CD19-mCherry plasmids with NEB PCR mutant kits. PCR-amplified constructs including mCherry, CD19-eGFP-Re#1, CD19-mCherry-Re#1 and CD19-ΔIC-mCherry-Re#1 from pCDNA3.1 were cloned into pLENTi-TRC313-Hgro vector, respectively.

To generate KPNA1-RE#1 cDNA resistant to KPNA1 sgRNA#1 sequence, the pDONR221-KPNA1 (HsCD00042508, DNASU) was modified to introduce the KPNA1 sgRNA #1 PAM sequence using NEB PCR mutant kit. The modified KPNA1 was then subcloned into pLENTi-TRC313-Hygro vector using Gateway cloning.

### Human genome-wide CRISPR screens and analysis

Daudi-Cas9 cells were used for the genome-wide CRISPR/Cas9 screen to identify essential genes for EBV entry into B cells. The Brunello library, which contains sgRNAs targeting 19,114 host genes, was obtained from Broad Institute. For each screen replicate, 150 million Cas9-expressing Daudi cells were infected with Brunello lentiviruses at a multiplicity of infection (MOI) of 0.3, to minimize lentivirus coinfection. Infection was performed in four 12-well plates per replicate. Cells were spun at 300g for 2 hours at 30°C in the presence of 4 µg/ml polybrene to increase transduction efficiency. After infection, the plates were incubated at 37°C for an additional 6 hours before reseeding into four T175 flasks. 48 hours post-infection, cells were selected with puromycin at 2 µg/ml. After six days of puromycin selection, 80 million cells were harvested and stored at -80°C as input. Simultaneously, 80 million cells were infected with EBV at a MOI of 10 for 2 hours, after which 300 mL RPMI 1640 medium (GIBCO) supplemented with 10% fetal bovine serum (FBS, GIBCO), 1% streptomycin and 1% penicillin were added. After 72 hours of culture, cells were harvested and sorted about 3 - 5% eGFP lower populations using FACS at the Brigham and Women’s Hospital Human Immunology Center. Genomic DNA of sorted cells was extracted with Qiagen Blood and Cell Culture DNA Mini Kit and sent together with the input DNA for PCR amplification and next-generation sequencing to Broad Institute. The STARS algorithm was used to identify statistically significant screen hits.

### CRISPR gene knockout and analysis

For hit validation, individual gene CRISPR knockouts were performed using single sgRNAs from the Brunello library. Briefly, sgRNAs targeting the genes of interest were synthesized, annealed, and cloned into the pLentiGuide-Puro vector according to protocols from the Broad Institute. The cloned sgRNAs were then packaged into lentiviruses and filtered through a 0.45-um filter before transducing the target cells. Forty-eight hours post-infection, the medium was replaced with fresh medium containing 2 µg/ml puromycin to select for successfully transduced cells, and this selection was maintained for 2 days. Following selection, cells were allowed to grow for an additional 6 days. Knockout efficiency was then assessed by Western blotting and the impact on EBV infection was evaluated through EBV infection assays.

### cDNA rescue

Daudi or Akata (-) cells single cell clone CD19 or KPNA1 KO were subjected to lentiviral transduction with rescue vectors: pLENTi-TRC313-mCherry, pLENTi-TRC313-CD19-mCherry-RE#1, pLENTi-TRC313-CD19-ΔIC-mCherry-RE#1 or pLENTi-TRC313-KNPA1-RE#1. The constructs included CD19-RE#1 and KPNA1-RE#1 cDNAs, which contain a PAM site mutation that abrogated CRISPR editing. These PAM mutations confer resistance to CD19 #1 sgRNA or KPNA1#1 sgRNA sequence, respectively. Following transduction, cells were selected with hygromycin for 10 days. CD19 or KPNA1 expression was confirmed by FACS or western blot analysis, as indicated.

### Virus preparation and EBV labelling with EdC

To prepare 100x EBV or 100x EdC-labelled EBV, Akata (EBV+eGFP) cells were resuspended in FBS-free RPMI 1640 medium at a concentration of 2 x10^6^ cells/ml and induced with 0.2% (v/v) anti-human immunoglobulin G for 6 hours at 37°C. Following induction, the cells were cultured in fresh RPMI1640 medium supplemented with 4% FBS or supplemented with additional 6uM EdC for 3 days. Virus-containing supernatant was collected under sterile conditions, filtered through a 0.45 µm filter, concentrated 100-fold by high-speed centrifugation at 50,000 g for 2 hours, and then resuspended in fresh FBS-free RPMI1640 medium. Prepared virus was stored at - 80°C and thawed immediately before use for infection.

### Determination of MOI and Quantification of EBV by Green Daudi Units (GDU)

Determination of MOI and quantification of EBV by GDUs was performed as described(40) . Briefly, Daudi cells were exposed to EBV-eGFP virus stocks for 2 hours at 37°C. Subsequently, 800 µl of fresh RPMI1640 medium supplemented with 10% FBS was added. 3 days post-infection, the percentage of eGFP-positive cells (EBV-infected cells) was determined using flow cytometry. The median infectious dose (ID50), defined as the virus dilution that results in 50% of the Daudi cells being eGFP-positive, was employed to estimate the virus concentration in GDUs per milliliter (mL). Typically, EBV stocks contain 1 to 2 x10^8^ GDUs/ml in 1 mL of a 100xEBV stocks solution.

### EBV infection of B cells

Daudi or EBV-Akata cells (0.5 or 1 x 10^6^) in 100 µl of RPMI1640 were co-incubated with 100 µl of 100x EBV for 2-3 hours at 37 °C. After co-incubation, unbound virus was removed by one time washing the cells with 5ml RPMI1640 medium. Infected cells were cultured in fresh medium for an additional 24 to 72 hours, as indicated. eGFP positive cell percentage was determined using FACS or fluorescence microscopy.

### EBV labelling with Alexa Fluor 594

Labelling of 200x EBV with Alexa Fluro 594 (Aime-Reactive probes; MP) was performed according to the published procedures(82) . Briefly, 4 mL of 200-fold concentrated EBV in FBS-free RPMI1640 was mixed with 400µl of 1mM Alexa Fluor 594, which was dissolved in dimethyl sulfoxide, in 0.2 M sodium bicarbonate buffer (pH 8.5). The mixture was gently rotated in the dark at room temperature for 1 hour. To separate the labelled EBV from the free dye, the mixture was purified using 25ml of RPMI1640 equilibrated PD-10 column (2.5ml/column) according to the manufacturer’s instructions. The first 1.5ml of flow-through was discarded, and the next 3 mL (1ml per tube) of the labeled virus fraction was collected. The labeled EBV was then aliquoted and stored at -80°C.

### Cell transfection and Immunoprecipitation

293T cells were transfected with the indicated plasmids with Lipofectamine 2000 (Invitrogen) according to the manufacturer’s instructions. 48 hours post-transfection, cells were lysed in a lysis buffer containing 50 mM Tris-HCl (pH 7.4), 250 mM NaCl, 5 mM EDTA (pH 8.0), 2.5mM MgCl2, 5% glycerol; 0.5% Nonidet P-40 (NP-40), 0.5% DOC and 0.5% Triton X-100, supplemented with 1 mM phenylmethylsulfonyl fluoride and Roche Complete protease inhibitor cocktail (04693159001, Roche). The lysates were centrifuged at 15,000g for 20 minutes at 4°C, and the supernatant was pre-cleared with 4 µg of mouse IgG and 25 µl of protein A/G magnetic beads for 1 - 2 hours at 4 °C. The pre-cleared lysate was then incubated with 25 μl of anti-HA magnetic beads (Invitrogen) for 8 hours at 4°C. Following five washes with lysis buffer to remove unbound proteins, the beads were resuspended in 1.25x SDS sample buffer and boiled for 5 minutes at 95°C. The complex was analyzed by western blotting using the indicated antibodies.

### Western blot

Western blotting analysis was performed as previously described. Briefly, cells were lysed in RIPA buffer containing a protease inhibitor mixture (Roche) and incubated on a rocker at 4°C for 15 minutes. The proteins were then separated by 10% SDS-PAGE and transferred to a nitrocellulose (NC) membrane, which was then probed with the indicated primary antibodies, followed by incubation with species-specific HRP-conjugated secondary antibodies. Immunoreactive bands were visualized using enhanced chemiluminescence (ECL, Pierce). To ensure equal protein loading, the same membranes were stripped and re-probed with mouse monoclonal antibodies against GAPDH.

### Immunofluorescence confocal microscopy

Akata (-)-CD19-eGFP cells were exposed to Alexa fluor 594-labelled EBV for 2 hours at 4°C, After two washes, the cells were visualized under a confocal fluorescence microscope.

Daudi Cells transfected with sgCTL or sgKPNA1#1 were exposed to AF594 or EdC-labelled EBV for 3 or 6 hours at 37°C. Cells were washed twice and fixed with 4% paraformaldehyde in PBS for 10 minutes. The cells were then washed twice with PBS, plated at 5 µl per well on 8-well slides, and allowed to dry and adhere for 30minutes. They were treated with 0.2% Triton-X100 in PBS for 5minutes, washed twice with PBS, and mounted using ProLong Diamond Antifade. For EdC staining, EdC-labelled EBV infected cells was processed by cycloaddition with azide-linked biotin and stained with avidin-AF647. Confocal images were acquired using a Zeiss confocal laser scanning microscope.

### Statistical analyses

P values were <0.05 were considered statistically significant, mean with ± SEM, as determined using Prism Version 10 (GraphPad). Data analysis was performed using t-test, depending on the data distribution and the number of comparison groups.

## LEAD CONTACT AND MATERIALS AVAILABILITY

Further information and requests for resources and reagents may be directed to Bo Zhao and Benjamin Gewurz (Lead Contact).

## AUTHOR CONTRIBUTIONS

Conceptualization, B.E.G., and B.Z.

Methodology, H.W., C.W. R.G.

Validation, H.W., C.W., D.Z., A.C. S.W., Y.S., S.Y., Y.Y., Y.L. G.H., and A.L.

Formal Analysis, R.G.

Resources, Z.W., S.J., B.E.G., and B.Z.

Writing – Original Draft, H.W., B.E.G., and B.Z.

Writing – Review & Editing, H.W., B.E.G., and B.Z.

Supervision B.E.G. and B.Z.

Funding Acquisition, B.E.G. and B.Z.

All authors approved the manuscript.

## ACKNOWLEDGMENTS

This work was funded by NIAID AI123420, AI192659 (B.Z.) and NIDCR R01DE033907 and NCI U01 R01CA275301 and P01CA269043 (B.E.G), by American Cancer Society Postdoctoral Fellowship PF-24-1308318-01-TBE (S.W.) and by K99DE034830 (Y.L). The authors gratefully acknowledge George and Sandra K. Schussel for their generous philanthropic support, which has been instrumental in advancing this work and furthering our mission to improve patient care through innovation and discovery.

## References

1. Cohen JI, Fauci AS, Varmus H, Nabel GJ. 2011. Epstein-Barr virus: an important vaccine target for cancer prevention. Sci Transl Med 3:107fs7.

2. Longnecker R KE, Cohen JI. 2013. Epstein-Barr Virus, 8th ed, vol Vol 2. Lippincott Williams & Wilkins, a Wolters Kluwer Business . Philadelphia.

3. Winter JR, Jackson C, Lewis JE, Taylor GS, Thomas OG, Stagg HR. 2020. Predictors of Epstein-Barr virus serostatus and implications for vaccine policy: A systematic review of the literature. J Glob Health 10:010404.

4. Wong Y, Meehan MT, Burrows SR, Doolan DL, Miles JJ. 2022. Estimating the global burden of Epstein-Barr virus-related cancers. J Cancer Res Clin Oncol 148:31–46.

5. Bjornevik K, Cortese M, Healy BC, Kuhle J, Mina MJ, Leng Y, Elledge SJ, Niebuhr DW, Scher AI, Munger KL, Ascherio A. 2022. Longitudinal analysis reveals high prevalence of Epstein-Barr virus associated with multiple sclerosis. Science 375:296–301.

6. Shoham T, Rajapaksa R, Kuo CC, Haimovich J, Levy S. 2006. Building of the tetraspanin web: distinct structural domains of CD81 function in different cellular compartments. Mol Cell Biol 26:1373–85.

7. Ojala PM, Sodeik B, Ebersold MW, Kutay U, Helenius A. 2000. Herpes simplex virus type 1 entry into host cells: reconstitution of capsid binding and uncoating at the nuclear pore complex in vitro. Mol Cell Biol 20:4922–31.

8. Pasdeloup D, Blondel D, Isidro AL, Rixon FJ. 2009. Herpesvirus capsid association with the nuclear pore complex and viral DNA release involve the nucleoporin CAN/Nup214 and the capsid protein pUL25. J Virol 83:6610–23.

9. Sodeik B, Ebersold MW, Helenius A. 1997. Microtubule-mediated transport of incoming herpes simplex virus 1 capsids to the nucleus. J Cell Biol 136:1007–21.

10. Copeland AM, Newcomb WW, Brown JC. 2009. Herpes simplex virus replication: roles of viral proteins and nucleoporins in capsid-nucleus attachment. J Virol 83:1660–8.

11. Younis S, Moutusy SI, Rasouli S, Jahanbani S, Pandit M, Wu X, Acharya S, Sharpe O, Wijeratne TU, Harris ML, Yang EY, Chaichian Y, Parsafar S, Baker MC, Harley JB, Meffre E, Steinman L, Marshak-Rothstein A, James JA, Martinez OM, Utz PJ, Orange DE, Lanz TV, Robinson WH. 2025. Epstein-Barr virus reprograms autoreactive B cells as antigen-presenting cells in systemic lupus erythematosus. Sci Transl Med 17:eady0210.

12. Laurynenka V, Harley JB. 2024. The 330 risk loci known for systemic lupus erythematosus (SLE): a review. Front Lupus 2.

13. Robinson WH, Younis S, Love ZZ, Steinman L, Lanz TV. 2024. Epstein-Barr virus as a potentiator of autoimmune diseases. Nat Rev Rheumatol 20:729–740.

14. Harley JB, Chen X, Pujato M, Miller D, Maddox A, Forney C, Magnusen AF, Lynch A, Chetal K, Yukawa M, Barski A, Salomonis N, Kaufman KM, Kottyan LC, Weirauch MT. 2018. Transcription factors operate across disease loci, with EBNA2 implicated in autoimmunity. Nat Genet 50:699–707.

15. Hong T, Parameswaran S, Donmez OA, Miller D, Forney C, Lape M, Saint Just Ribeiro M, Liang J, Edsall LE, Magnusen AF, Miller W, Chepelev I, Harley JB, Zhao B, Kottyan LC, Weirauch MT. 2021. Epstein-Barr virus nuclear antigen 2 extensively rewires the human chromatin landscape at autoimmune risk loci. Genome Res 31:2185–2198.

16. Hutt-Fletcher LM. 2007. Epstein-Barr virus entry. J Virol 81:7825–32.

17. Bu GL, Xie C, Kang YF, Zeng MS, Sun C. 2022. How EBV Infects: The Tropism and Underlying Molecular Mechanism for Viral Infection. Viruses 14.

18. Shannon-Lowe C, Rowe M. 2014. Epstein Barr virus entry; kissing and conjugation. Curr Opin Virol 4:78–84.

19. Tanner J, Weis J, Fearon D, Whang Y, Kieff E. 1987. Epstein-Barr virus gp350/220 binding to the B lymphocyte C3d receptor mediates adsorption, capping, and endocytosis. Cell 50:203–13.

20. Ogembo JG, Kannan L, Ghiran I, Nicholson-Weller A, Finberg RW, Tsokos GC, Fingeroth JD. 2013. Human complement receptor type 1/CD35 is an Epstein-Barr Virus receptor. Cell Rep 3:371–85.

21. Escalante GM, Mutsvunguma LZ, Muniraju M, Rodriguez E, Ogembo JG. 2022. Four Decades of Prophylactic EBV Vaccine Research: A Systematic Review and Historical Perspective. Front Immunol 13:867918.

22. Li Q, Spriggs MK, Kovats S, Turk SM, Comeau MR, Nepom B, Hutt-Fletcher LM. 1997. Epstein-Barr virus uses HLA class II as a cofactor for infection of B lymphocytes. J Virol 71:4657–62.

23. Wang X, Hutt-Fletcher LM. 1998. Epstein-Barr virus lacking glycoprotein gp42 can bind to B cells but is not able to infect. J Virol 72:158–63.

24. Haan KM, Kwok WW, Longnecker R, Speck P. 2000. Epstein-Barr virus entry utilizing HLA-DP or HLA-DQ as a coreceptor. J Virol 74:2451–4.

25. Haan KM, Longnecker R. 2000. Coreceptor restriction within the HLA-DQ locus for Epstein-Barr virus infection. Proc Natl Acad Sci U S A 97:9252–7.

26. Hardy S, Chhan CB, Davis AR, McGuire AT. 2025. Viral Entry. Curr Top Microbiol Immunol doi:10.1007/82_2025_300.

27. Wang X, Kenyon WJ, Li Q, Mullberg J, Hutt-Fletcher LM. 1998. Epstein-Barr virus uses different complexes of glycoproteins gH and gL to infect B lymphocytes and epithelial cells. J Virol 72:5552–8.

28. Molesworth SJ, Lake CM, Borza CM, Turk SM, Hutt-Fletcher LM. 2000. Epstein-Barr virus gH is essential for penetration of B cells but also plays a role in attachment of virus to epithelial cells. J Virol 74:6324–32.

29. Silva AL, Omerovic J, Jardetzky TS, Longnecker R. 2004. Mutational analyses of Epstein-Barr virus glycoprotein 42 reveal functional domains not involved in receptor binding but required for membrane fusion. J Virol 78:5946–56.

30. Wu L, Borza CM, Hutt-Fletcher LM. 2005. Mutations of Epstein-Barr virus gH that are differentially able to support fusion with B cells or epithelial cells. J Virol 79:10923–30.

31. Sathiyamoorthy K, Hu YX, Mohl BS, Chen J, Longnecker R, Jardetzky TS. 2016. Structural basis for Epstein-Barr virus host cell tropism mediated by gp42 and gHgL entry glycoproteins. Nat Commun 7:13557.

32. Sathiyamoorthy K, Jiang J, Mohl BS, Chen J, Zhou ZH, Longnecker R, Jardetzky TS. 2017. Inhibition of EBV-mediated membrane fusion by anti-gHgL antibodies. Proc Natl Acad Sci U S A 114:E8703–E8710.

33. Mohl BS, Chen J, Sathiyamoorthy K, Jardetzky TS, Longnecker R. 2016. Structural and Mechanistic Insights into the Tropism of Epstein-Barr Virus. Mol Cells 39:286–91.

34. Sathiyamoorthy K, Jiang J, Hu YX, Rowe CL, Mohl BS, Chen J, Jiang W, Mellins ED, Longnecker R, Zhou ZH, Jardetzky TS. 2014. Assembly and architecture of the EBV B cell entry triggering complex. PLoS Pathog 10:e1004309.

35. Bu W, Kumar A, Board NL, Kim J, Dowdell K, Zhang S, Lei Y, Hostal A, Krogmann T, Wang Y, Pittaluga S, Marcotrigiano J, Cohen JI. 2024. Epstein-Barr virus gp42 antibodies reveal sites of vulnerability for receptor binding and fusion to B cells. Immunity 57:559–573 e6.

36. Li Y, Zhang H, Sun C, Dong XD, Xie C, Liu YT, Lin RB, Kong XW, Hu ZL, Ma XY, Dai DL, Zhu QY, Li YC, Li Y, Liu SX, Yuan L, Zhou PH, Gao S, Tang YP, Yang JY, Han P, McGuire AT, Zhao B, Bei JX, Robertson E, Zeng YX, Zhong Q, Zeng MS. 2025. R9AP is a common receptor for EBV infection in epithelial cells and B cells. Nature 644:205–213.

37. Wang HB, Zhang H, Zhang JP, Li Y, Zhao B, Feng GK, Du Y, Xiong D, Zhong Q, Liu WL, Du H, Li MZ, Huang WL, Tsao SW, Hutt-Fletcher L, Zeng YX, Kieff E, Zeng MS. 2015. Neuropilin 1 is an entry factor that promotes EBV infection of nasopharyngeal epithelial cells. Nat Commun 6:6240.

38. Zhang H, Li Y, Wang HB, Zhang A, Chen ML, Fang ZX, Dong XD, Li SB, Du Y, Xiong D, He JY, Li MZ, Liu YM, Zhou AJ, Zhong Q, Zeng YX, Kieff E, Zhang Z, Gewurz BE, Zhao B, Zeng MS. 2018. Ephrin receptor A2 is an epithelial cell receptor for Epstein-Barr virus entry. Nat Microbiol 3:1–8.

39. Zhang H, Li YC, Pang D, Xie C, Zhang T, Li Y, Li Y, Jiang ZY, Bu GL, Liu MM, Chen YR, Fei HX, Lin RB, Wu PH, Du WT, Zhao GX, Luo YL, Han P, Zhong Q, Sun C, Zeng MS. 2025. Desmocollin 2 is a dominant entry receptor for Epstein-Barr virus infection of epithelial cells. Nat Microbiol doi:10.1038/s41564-025-02067-8.

40. Wang H, Mou Z, Yeo YY, Ge Q, Liu X, Narita Y, Li Z, Wang C, Li W, Zhao KR, Li J, Bu W, Gewurz B, Cohen JI, Teng M, Dai X, Liu X, Jiang S, Zhao B. 2025. Epstein-Barr virus exploits desmocollin 2 as the principal epithelial cell entry receptor. Nat Microbiol doi:10.1038/s41564-025-02126-0.

41. Chen J, Sathiyamoorthy K, Zhang X, Schaller S, Perez White BE, Jardetzky TS, Longnecker R. 2018. Ephrin receptor A2 is a functional entry receptor for Epstein-Barr virus. Nat Microbiol 3:172–180.

42. Chesnokova LS, Nishimura SL, Hutt-Fletcher LM. 2009. Fusion of epithelial cells by Epstein-Barr virus proteins is triggered by binding of viral glycoproteins gHgL to integrins alphavbeta6 or alphavbeta8. Proc Natl Acad Sci U S A 106:20464–9.

43. Mohl BS, Chen J, Longnecker R. 2019. Gammaherpesvirus entry and fusion: A tale how two human pathogenic viruses enter their host cells. Adv Virus Res 104:313–343.

44. Chesnokova LS, Jiang R, Hutt-Fletcher LM. 2015. Viral Entry. Curr Top Microbiol Immunol 391:221–35.

45. Chandran B, Hutt-Fletcher L. 2007. Gammaherpesviruses entry and early events during infection. In Arvin A, Campadelli-Fiume G, Mocarski E, Moore PS, Roizman B, Whitley R, Yamanishi K (ed), Human Herpesviruses: Biology, Therapy, and Immunoprophylaxis, Cambridge.

46. Torne AS, Robertson ES. 2024. Epigenetic Mechanisms in Latent Epstein-Barr Virus Infection and Associated Cancers. Cancers (Basel) 16.

47. Lieberman PM, Tempera I. 2025. Chromatin Control of EBV Infection and Latency. Curr Top Microbiol Immunol doi:10.1007/82_2025_318.

48. Munz C. 2025. Epstein-Barr virus pathogenesis and emerging control strategies. Nat Rev Microbiol 23:667–679.

49. Guo R, Gewurz BE. 2022. Epigenetic control of the Epstein-Barr lifecycle. Curr Opin Virol 52:78–88.

50. Hurley EA, Thorley-Lawson DA. 1988. B cell activation and the establishment of Epstein-Barr virus latency. J Exp Med 168:2059–75.

51. Pich D, Mrozek-Gorska P, Bouvet M, Sugimoto A, Akidil E, Grundhoff A, Hamperl S, Ling PD, Hammerschmidt W. 2019. First Days in the Life of Naive Human B Lymphocytes Infected with Epstein-Barr Virus. mBio 10.

52. Doench JG, Hartenian E, Graham DB, Tothova Z, Hegde M, Smith I, Sullender M, Ebert BL, Xavier RJ, Root DE. 2014. Rational design of highly active sgRNAs for CRISPR-Cas9-mediated gene inactivation. Nat Biotechnol 32:1262–7.

53. Tuveson DA, Carter RH, Soltoff SP, Fearon DT. 1993. CD19 of B cells as a surrogate kinase insert region to bind phosphatidylinositol 3-kinase. Science 260:986–9.

54. Degn SE, Tolar P. 2025. Towards a unifying model for B-cell receptor triggering. Nat Rev Immunol 25:77–91.

55. Susa KJ, Bradshaw GA, Eisert RJ, Schilling CM, Kalocsay M, Blacklow SC, Kruse AC. 2024. A spatiotemporal map of co-receptor signaling networks underlying B cell activation. Cell Rep 43:114332.

56. Wang K, Wei G, Liu D. 2012. CD19: a biomarker for B cell development, lymphoma diagnosis and therapy. Exp Hematol Oncol 1:36.

57. Tedder TF, Inaoki M, Sato S. 1997. The CD19-CD21 complex regulates signal transduction thresholds governing humoral immunity and autoimmunity. Immunity 6:107–18.

58. Susa KJ, Seegar TC, Blacklow SC, Kruse AC. 2020. A dynamic interaction between CD19 and the tetraspanin CD81 controls B cell co-receptor trafficking. Elife 9.

59. Maecker HT, Levy S. 1997. Normal lymphocyte development but delayed humoral immune response in CD81-null mice. J Exp Med 185:1505–10.

60. Shimizu N, Tanabe-Tochikura A, Kuroiwa Y, Takada K. 1994. Isolation of Epstein-Barr virus (EBV)-negative cell clones from the EBV-positive Burkitt’s lymphoma (BL) line Akata: malignant phenotypes of BL cells are dependent on EBV. J Virol 68:6069–73.

61. Martin H, Murray C, Christeller J, McGhie T. 2008. A fluorescence polarization assay to quantify biotin and biotin-binding proteins in whole plant extracts using Alexa-Fluor 594 biocytin. Anal Biochem 381:107–12.

62. Harris HJ, Farquhar MJ, Mee CJ, Davis C, Reynolds GM, Jennings A, Hu K, Yuan F, Deng H, Hubscher SG, Han JH, Balfe P, McKeating JA. 2008. CD81 and claudin 1 coreceptor association: role in hepatitis C virus entry. J Virol 82:5007–20.

63. Piboonniyom SO, Duensing S, Swilling NW, Hasskarl J, Hinds PW, Munger K. 2003. Abrogation of the retinoblastoma tumor suppressor checkpoint during keratinocyte immortalization is not sufficient for induction of centrosome-mediated genomic instability. Cancer Res 63:476–83.

64. Gao Y, Colletti K, Pari GS. 2008. Identification of human cytomegalovirus UL84 virus- and cell-encoded binding partners by using proteomics analysis. J Virol 82:96–104.

65. Khan H, Sumner RP, Rasaiyaah J, Tan CP, Rodriguez-Plata MT, Van Tulleken C, Fink D, Zuliani-Alvarez L, Thorne L, Stirling D, Milne RS, Towers GJ. 2020. HIV-1 Vpr antagonizes innate immune activation by targeting karyopherin-mediated NF-kappaB/IRF3 nuclear transport. Elife 9.

66. Schwarz TM, Edwards MR, Diederichs A, Alinger JB, Leung DW, Amarasinghe GK, Basler CF. 2017. VP24-Karyopherin Alpha Binding Affinities Differ between Ebolavirus Species, Influencing Interferon Inhibition and VP24 Stability. J Virol 91.

67. Boivin S, Hart DJ. 2011. Interaction of the influenza A virus polymerase PB2 C-terminal region with importin alpha isoforms provides insights into host adaptation and polymerase assembly. J Biol Chem 286:10439–48.

68. McBride KM, Banninger G, McDonald C, Reich NC. 2002. Regulated nuclear import of the STAT1 transcription factor by direct binding of importin-alpha. EMBO J 21:1754–63.

69. Reid SP, Leung LW, Hartman AL, Martinez O, Shaw ML, Carbonnelle C, Volchkov VE, Nichol ST, Basler CF. 2006. Ebola virus VP24 binds karyopherin alpha1 and blocks STAT1 nuclear accumulation. J Virol 80:5156–67.

70. Dohner K, Wolfstein A, Prank U, Echeverri C, Dujardin D, Vallee R, Sodeik B. 2002. Function of dynein and dynactin in herpes simplex virus capsid transport. Mol Biol Cell 13:2795–809.

71. Abaitua F, O’Hare P. 2008. Identification of a highly conserved, functional nuclear localization signal within the N-terminal region of herpes simplex virus type 1 VP1-2 tegument protein. J Virol 82:5234–44.

72. Abaitua F, Hollinshead M, Bolstad M, Crump CM, O’Hare P. 2012. A Nuclear localization signal in herpesvirus protein VP1-2 is essential for infection via capsid routing to the nuclear pore. J Virol 86:8998–9014.

73. Hennig T, Abaitua F, O’Hare P. 2014. Functional analysis of nuclear localization signals in VP1-2 homologues from all herpesvirus subfamilies. J Virol 88:5391–405.

74. Bauer DW, Huffman JB, Homa FL, Evilevitch A. 2013. Herpes virus genome, the pressure is on. J Am Chem Soc 135:11216–21.

75. Brandariz-Nunez A, Liu T, Du T, Evilevitch A. 2019. Pressure-driven release of viral genome into a host nucleus is a mechanism leading to herpes infection. Elife 8.

76. Dohner K, Ramos-Nascimento A, Bialy D, Anderson F, Hickford-Martinez A, Rother F, Koithan T, Rudolph K, Buch A, Prank U, Binz A, Hugel S, Lebbink RJ, Hoeben RC, Hartmann E, Bader M, Bauerfeind R, Sodeik B. 2018. Importin alpha1 is required for nuclear import of herpes simplex virus proteins and capsid assembly in fibroblasts and neurons. PLoS Pathog 14:e1006823.

77. Pumroy RA, Cingolani G. 2015. Diversification of importin-alpha isoforms in cellular trafficking and disease states. Biochem J 466:13–28.

78. Yasuhara N, Shibazaki N, Tanaka S, Nagai M, Kamikawa Y, Oe S, Asally M, Kamachi Y, Kondoh H, Yoneda Y. 2007. Triggering neural differentiation of ES cells by subtype switching of importin-alpha. Nat Cell Biol 9:72–9.

79. Farina A, Feederle R, Raffa S, Gonnella R, Santarelli R, Frati L, Angeloni A, Torrisi MR, Faggioni A, Delecluse HJ. 2005. BFRF1 of Epstein-Barr virus is essential for efficient primary viral envelopment and egress. J Virol 79:3703–12.

80. Gonnella R, Farina A, Santarelli R, Raffa S, Feederle R, Bei R, Granato M, Modesti A, Frati L, Delecluse HJ, Torrisi MR, Angeloni A, Faggioni A. 2005. Characterization and intracellular localization of the Epstein-Barr virus protein BFLF2: interactions with BFRF1 and with the nuclear lamina. J Virol 79:3713–27.

81. Yiu SPT, Liao Y, Yan J, Weekes MP, Gewurz BE. 2025. Epstein-Barr virus BALF0/1 subverts the Caveolin and ERAD pathways to target B cell receptor complexes for degradation. Proc Natl Acad Sci U S A 122:e2400167122.

82. Zhang S, Tan HC, Ooi EE. 2011. Visualizing dengue virus through Alexa Fluor labeling. J Vis Exp doi:10.3791/3168:e3168.

